# Recipient cell identity governs intracellular transport and membrane interactions of extracellular vesicles across species

**DOI:** 10.64898/2026.06.03.729830

**Authors:** Bo Have Brøchner, Ping Song, Yan Yan, Hannah Weissinger, Elisabeth Asta Sørensen, Jørgen Kjems, Mette Galsgaard Malle

**Author notes:** To whom correspondence should be addressed at (J.K) or (M.G.M).

## Abstract

Extracellular vesicles (EVs) function as a natural communication system, enabling transfer of biomolecules between cells. Although EVs are produced by nearly all cell types and have shown promise as therapeutic agents, our understanding of how EVs from different cellular origins recognize recipient cells, are taken up, and processed intracellularly remains limited. In particular, it remains unclear whether intracellular EV behavior is primarily governed by EV-intrinsic properties or by the recipient-cell environment. Here, we systematically investigate the relative contributions of EV origin and recipient-cell identity to intracellular EV trafficking. By combining single-vesicle tracking with machine learning-based diffusion-state classification, we analyzed EVs derived from four distinct sources across three recipient cell types over three different time spans. This approach enabled characterization of dynamic transport behaviors, overcoming limitations of traditional ensemble-averaging methods. We observed recipient cell-dependent differences in intracellular trafficking patterns, ranging from relatively uniform and dynamic motion to more heterogeneous and confined behaviors, reflecting variability in intracellular transport environments. Despite these differences, EV origin had only a modest influence on intracellular dynamics following uptake. These findings suggest a model in which intracellular transport arises from an interplay between general EV-associated properties and cell-specific environment.

## Introduction

Extracellular vesicles (EVs) are lipid bilayer-enclosed nanoparticles secreted by all cell types, and play a central role in intercellular communication through the transfer of biomolecules, including proteins, lipids, and nucleic acids (1-3). Owing to the intrinsic biocompatibility, low immunogenicity, and ability to cross biological barriers, EVs have emerged as a promising candidate for drug delivery and diagnostic applications (4-10). However, EVs are highly heterogeneous, and their biological function and targeting capacity are not fully characterized. In particular, the relative contributions of EV origin and recipient-cell identity to intracellular EV behavior remain poorly understood.

To understand the behavior, fluorescently labelled or tagged EVs have been used in combination with flow cytometry or microscopy, revealing various underlying mechanisms (11-13). Often, studies compare EVs from several origins, showing either no significant differences or preferential uptake, however only using a single cell type (11, 14-18), reflecting the long-standing debate in the field on whether EV origin has an impact on uptake. Other studies use the opposite approach, looking at how EVs from a specific cell type are internalized across several different recipient cell lines (19-21), showing that EVs can be taken up in all cell lines but can differ widely in the amount taken up.

Taken together, existing studies indicate that EV uptake is inherently heterogeneous, varying across both EV sources and recipient cell types. Recent studies have gone beyond qualitative uptake studies to get a detailed understanding of the dynamic behavior of EVs by unmasking the heterogeneity using single particle imaging and tracking analysis (6, 22, 23). However, interpreting these trajectories remains challenging due to the heterogeneous and dynamic nature of intracellular transport as well as the difficulty of obtaining high-quality data with sufficient temporal resolution. EVs can exhibit multiple modes of motion, including free diffusion, active transport along the cytoskeletal structures, confinement at membranes, and hindered movement in crowded intracellular environments. Most studies use the technique to state that it is possible to differentiate between EV types (7, 11, 24) or compare how EVs behave differently on different recipient cells (25). Traditional analytical approaches, such as mean squared displacement (MSD), often fail to resolve the dynamic transition as they average over the entire trajectory, masking the complex nature of intracellular transport. Recent machine learning-based trajectory segmentation improves motion-state classification, but it remains unclear whether intracellular EV behavior is driven by EV-intrinsic properties or the recipient cell environment (22, 24, 26).

To address this gap, we systematically investigated how EV origin and recipient-cell identity influence intracellular EV trafficking by combining single-vesicle tracking with single-particle diffusion-state classification. Using EVs derived from four sources and three recipient cell types, we aimed to define the relative contributions of vesicle origin and recipient-cell identity to EV uptake and subsequent intracellular processes. In doing so, we sought to establish whether intracellular EV trafficking is specified predominantly by EV-intrinsic properties or by their interplay with the intracellular transport environment of the recipient cell. Answering this question is important for understanding EV-mediated communication in complex biological systems and for guiding the development of EV-based therapeutic strategies.

## Results

### Extracellular vesicles from diverse biological sources

To investigate how the origin of extracellular vesicles (EVs) influences their behavior in recipient cells, EVs were isolated from four distinct sources: an immortalized mouse myoblast cell line, C2C12 (C2C12 EVs), human plasma (hP-EVs), mouse plasma (mP-EVs), and fresh bovine milk (bM-EVs). These sources represent EVs derived from cultured cells, circulating plasma, and dietary sources, spanning EVs originating from controlled in vitro systems, physiological circulation, and cross-species nutritional environments. EVs were subsequently applied to three recipient cell lines: bEnd.3, an immortalized mouse brain endothelial cell line derived from BALB/c mouse cerebral cortex, HEK293-H, a fast-growing variant of the human embryonic kidney cell line, and ARPE-19, an immortalized human retinal pigment epithelial cell line. By combining real-time quantitative microscopy with single-vesicle tracking in live cells, we investigated whether EV uptake and intracellular processing reflect intrinsic properties of EV origin, features of the recipient cell, or both (Figure 1).

**Figure 1:**
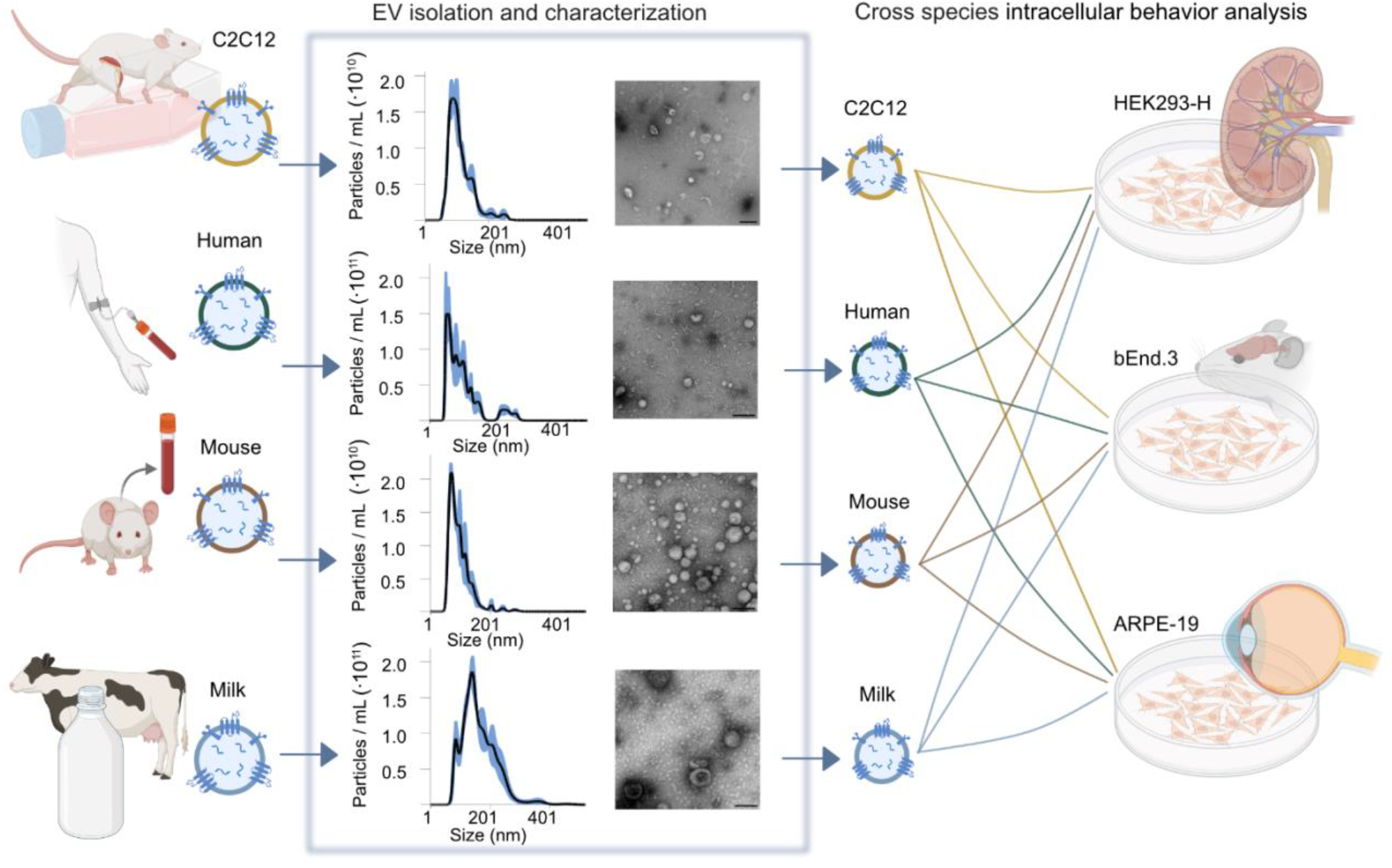
Multi-species extracellular vesicle isolation and characterization. A Schematic of the experimental approach. EVs derived from four different origins, C2C12 cells, human plasma, mouse plasma, and fresh bovine milk (left panel), were isolated and characterized by NTA and nsTEM (middle panel), then added to three different recipient cell lines (bEnd.3, HEK293-H, and ARPE-19; right panel) to understand EV uptake and intracellular behavior. Scale bars on nsTEM: 100 nm.

EVs were isolated using ultracentrifugation followed by size-exclusion chromatography (SEC) purification for bM-EVs and C2C12 EVs and using SEC for mP-EVs and hP-EVs (see methods), in accordance with the MISEV guidelines (27). Western blot analysis confirmed enrichment of the EV markers CD81 and flotillin-1 in the EV fraction, whereas the endoplasmic reticulum marker calnexin was predominantly detected in the cellular fraction (Supplementary Figure 1). Particle size and concentration were determined by nanoparticle tracking analysis (NTA). Mean particle diameters were 102.1 ± 1.0 nm, 91.0 ± 3.9 nm, 87.4 ± 1.0 nm, and 146.0 ± 3.5 nm for C2C12-, hP-, mP-, and bM EVs, respectively (Figure 1 and Supplementary Figure 2), consistent with the expected size range for small EVs and previous reports (28). EV morphology was examined using negative-stain transmission electron microscopy (nsTEM). All preparations displayed spherical to oval vesicular structures, consistent with expected EV morphology (Figure 1) (14, 29, 30).

**Figure 2:**
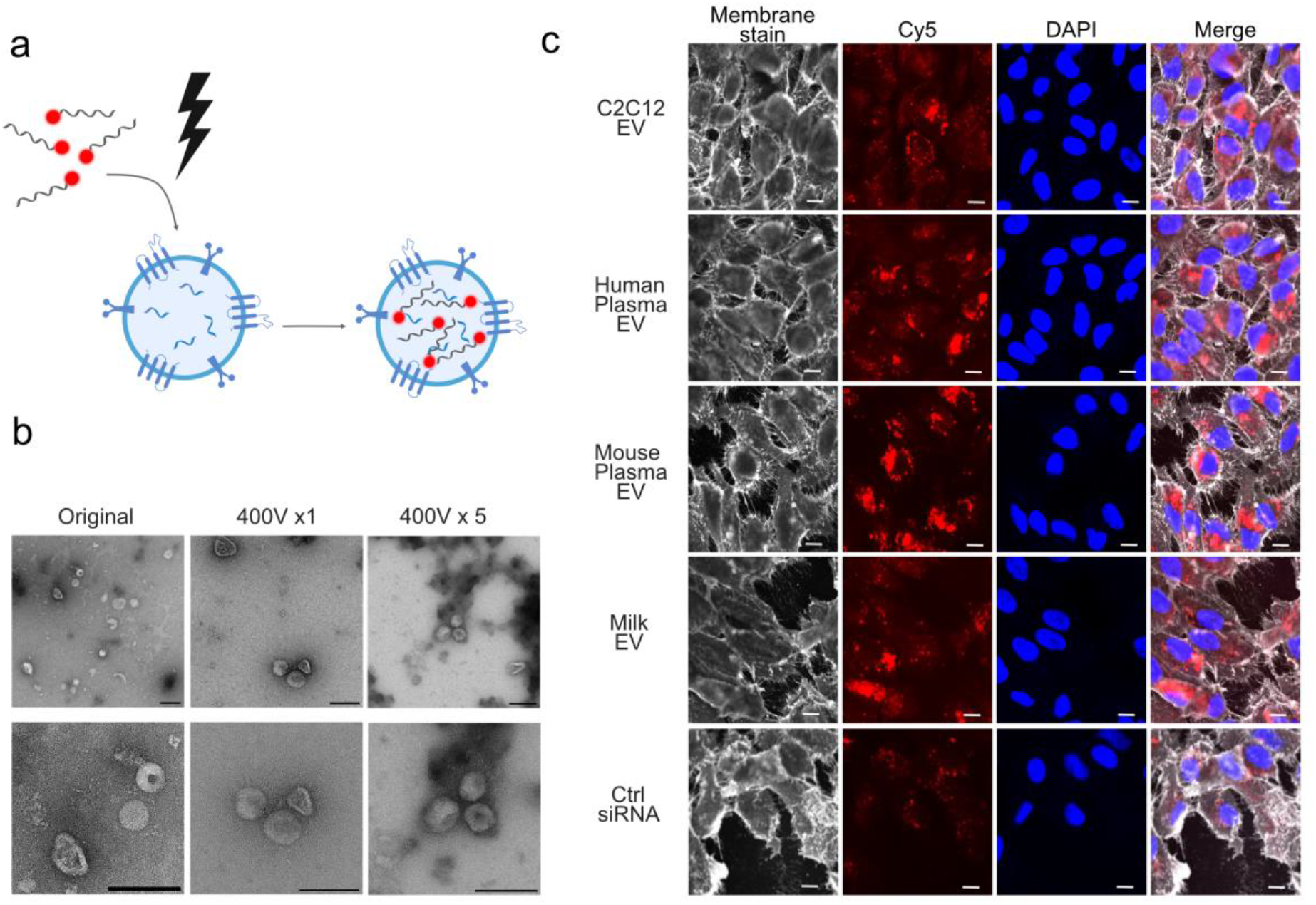
Single-vesicle quantification and optimized siRNA loading for EV-mediated RNA delivery. a) Schematic illustration of electroporation of Cy5-siRNA into EVs. b) Representative nsTEM images of EVs from C2C12 cells, showing vesicles with typical EV morphology and no obvious structural differences after either 400V 1x or 5x pulse electroporation. Scalebar 100 nm. The image of EVs before electroporation is reused from Figure 1. c) Cellular uptake of EV delivered Cy5-siRNA after 24h, showing representative delivery in ARPE-19 cells. Scalebar 10μm.

### Consistent Uptake and Cargo Delivery Across EV Sources and Recipient Cell Types

To assess the impact of EV origin on cargo delivery to different recipient cells, the EV membrane was labelled with ATTO 488-DOPE, and Cy5-labeled siRNA was electroporated into lumen (Figure 2a), enabling co-visualization of EV membrane and intracellular cargo delivery. First, encapsulation efficiency of Cy5-labled siRNA was quantified at the single-vesicle level using total internal reflection fluorescence (TIRF) microscopy using a colocalization assay (Supplementary Figure 3). Quantification was performed by detecting >155000 membrane-labeled EVs using an in-house developed Python software and determining the percentage of vesicles displaying colocalized Cy5-siRNA signal above background levels (Supplementary Figure 3 and Supplementary Figure 4).

**Figure 3:**
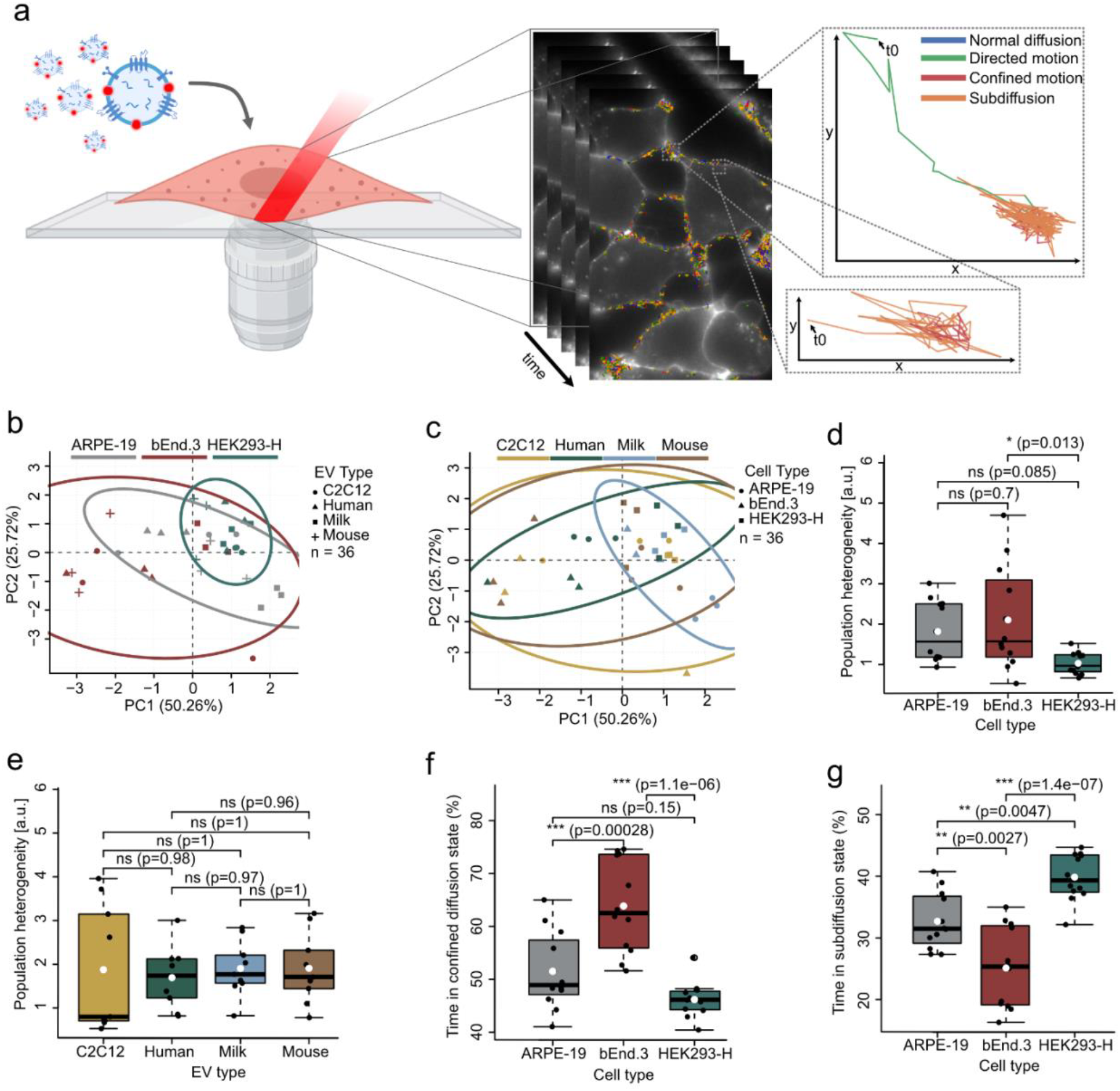
Multivariate analysis of diffusional behaviors of EVs in different recipient cell types. a) Segmentation of trajectories obtained by tracking single vesicles in live cells in real time. Zoom-in on two trajectories showing transition between diffusional states over time, underlining the complex behavior of particles inside cells. b,c) PCA score plot showing clustering of the amount of time spent in each state derived from single-particle trajectories across 36 HILO movies, grouped by imaged cell type (b) or EV origin (c). d,e) Comparison of within-group heterogeneity according to cell type (d) or EV origin (e). Individual datapoints are depicted with a boxplot showing the mean (white dot). f,g) Variables exhibiting the highest absolute loadings on PC1 and PC2, respectively, were selected as key contributors to component structure. Boxplots show the distribution of these variables across experimental groups. Asterisks (*, **, ***) in (d-g) indicate a *P ≤ 0.05, **P ≤ 0.01, and ***P ≤ 0.001 and reflect a significant difference assessed by one-way ANOVA with Tukey post hoc tests.

**Figure 4:**
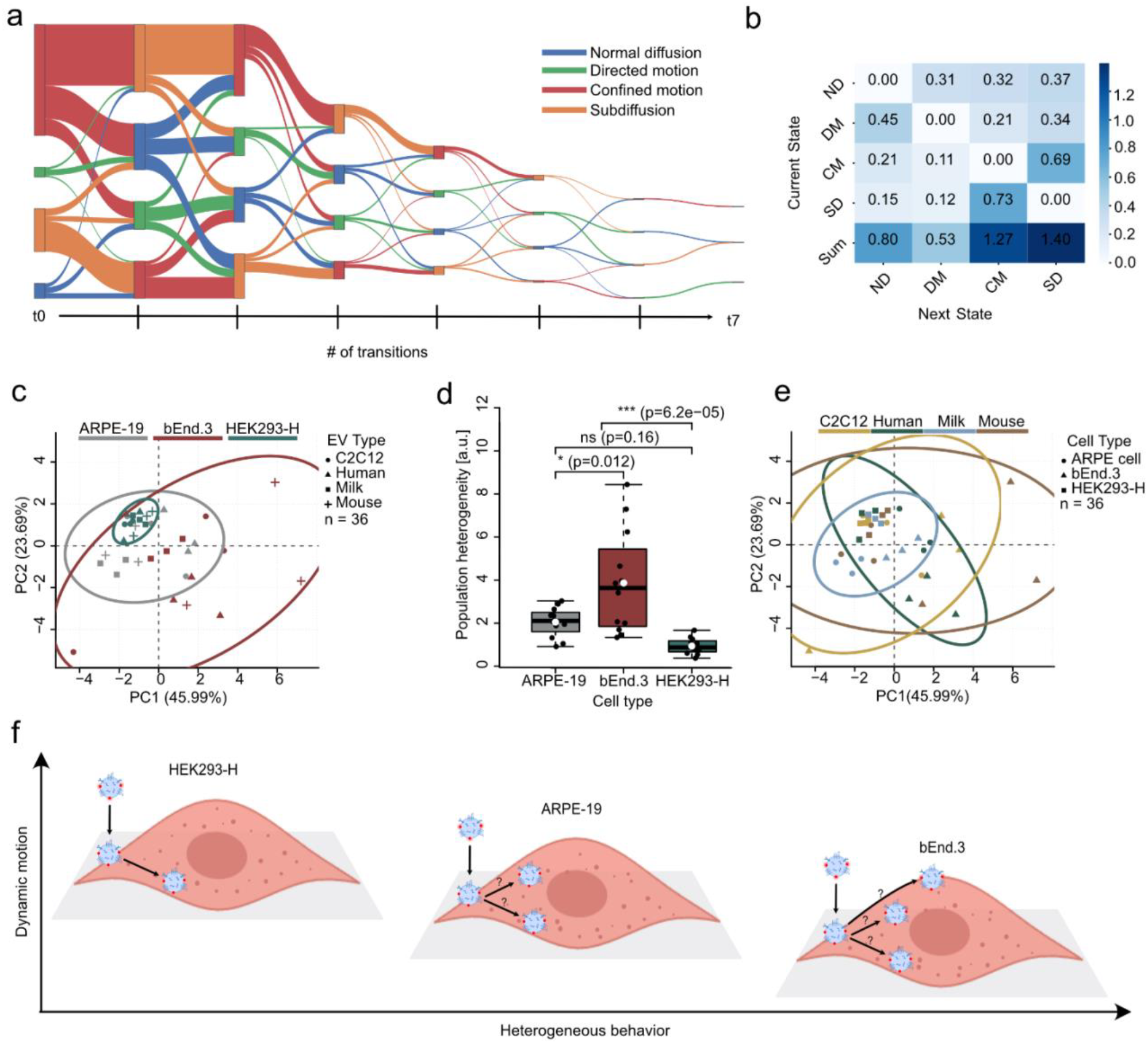
Multivariate analysis of transition probabilities of EVs in different recipient cell types. a) Sankey diagram of single-particle diffusive behavior over time, illustrating transitions between distinct diffusion modes along individual trajectories. The plot shows the fraction of particles transitioning between states from the initial state (t_0_) through the 7th transition, with later transitions represented by a subset of particles due to differences in the number of state transitions. Confined, subdiffusive, normal diffusive, and directed motion are shown in red, orange, blue, and green, respectively. b) Transition Probability Matrix. c) PCA plot showing clustering of transition probabilities derived from single-particle trajectories across 36 HILO movies, grouped by imaged cell type. d) Comparison of within-group heterogeneity according to cell type. Individual datapoints are depicted with a boxplot showing the mean (white dot). c) PCA plot showing clustering of transition probabilities derived from single-particle trajectories across 36 HILO movies, grouped by imaged EV origin. f) Schematic illustration of diffusional behavior in HEK293-H, ARPE-19, and bEnd.3 cells, showing heterogeneous behavior of EVs in cells on the x-axis and the amount of dynamic motion on the y-axis. a and b are representative data from Human plasma EVs in HEK-293-H cells. Asterisks (*, **, ***) in (e) indicate a *P ≤ 0.05, **P ≤ 0.01, and ***P ≤ 0.001 and reflect a significant difference assessed by one-way ANOVA with Tukey post hoc tests.

To evaluate whether the electroporation procedure affected vesicle integrity, EVs were examined by nsTEM before and after loading. Vesicles retained a spherical morphology under both single-and five-pulse 400V electroporation conditions, indicating preserved structural integrity (Figure 2b). However, increasing voltage to 750V led to aggregation (Supplementary Figure 3c).

Next, EV uptake and cargo delivery were assessed in recipient cells. Here, cells were incubated with EVs containing encapsulated Cy5-siRNA and analyzed by confocal microscopy after both 4 h and 24 h for subsequently washed and fixed cells. Clear intracellular Cy5 fluorescence was observed in all recipient cell types, indicating cellular uptake and cytoplasmic delivery of siRNA from all EV origins - see Figure 2c for 24 h experiment in ARPE cells. See Supplementary Figure 5-7 for additional time points and recipient cell types. Electroporated Cy5-siRNA alone produced a distinct pattern (ctrl siRNA; Figure 2c and Supplementary Figure 7), supporting that fluorescence in Cy5-siRNA-loaded EVs arises from EV-mediated delivery rather than siRNA aggregates (31).

Collectively, confocal microscopy uptake analysis showed successful intracellular siRNA delivery with comparable signal delivery across all EV origins and recipient cell types, suggesting that EV interaction and content delivery efficiency are not strongly dependent on EV source or recipient cell type.

### Recipient cell type determines intracellular EV diffusional behavior

To investigate how EVs are trafficked after cellular uptake, we quantified the diffusional behavior of individual EVs using single-particle tracking across the four EV sources and three recipient cell types.

Here, live-cell imaging was performed using highly inclined and laminated optical sheet (HILO) microscopy, which enables high signal-to-noise detection of fluorescent EVs both at the plasma membrane and within the cytoplasm. Individual diffraction-limited EVs were localized and linked into trajectories describing their motion inside cells (Figure 3a and Supplementary Figure 8a. In total, >440,000 EV trajectories were analyzed using in-house developed Python software. Conventional diffusion coefficients estimated from full trajectory mean squared displacement (MSD) analysis did not resolve differences by EV origin or recipient cell type (Supplementary Fig. 8b). Notably, MSD fits were poor for trajectories exhibiting transitions between different diffusional states (Supplementary Figure 8c), highlighting a fundamental limitation of full-trajectory MSD analysis when applied to temporally heterogeneous motion (26). To more accurately capture the complex spatiotemporal behavior of EVs inside cells, trajectories were therefore segmented using a recently developed deep-learning-based model, enabling classification into four biological-behavioral descriptive states: normal diffusion, directed motion, confined diffusion, and subdiffusive motion, reflecting respectively free diffusion, active transport along the cytoskeletal structures, confinement at membranes, and hindered movement in crowded intracellular environments (Figure 3a) (26).

Using this framework, we quantified the fraction of time each EV occupied diffusional states to identify initial cell interaction and subsequent trafficking (Supplementary Figure 9). Interestingly, we found that EV size does not influence diffusional behavior when using membrane signal intensity as a proxy for EV size (16, 24), with no size-dependent preference for any diffusion state observed (Supplementary Figure 10-12), consistent with reports showing similar intracellular dynamics across distinct EV size fractions (24).

Principal component analysis (PCA) of diffusional state distributions revealed partial separation of EV spatiotemporal trajectories according to recipient cell type (Figure 3b). EVs tracked in either ARPE-19, bEnd.3, and HEK293-H cells occupied partially distinct regions of the principal component space, revealing that recipient cell identity is a major determinant of intracellular EV diffusional behavior. In contrast, PCA grouped by EV origin showed substantial overlap between EVs derived from C2C12 cells, human plasma, mouse plasma, and milk (Figure 3c), independent of the recipient cell, suggesting that EV origin has a comparatively minor influence on intracellular trafficking dynamics.

Analysis of heterogeneity within each recipient cell type further supported this observation. EV trajectories from the four distinct EV sources in HEK293-H cells displayed significantly lower heterogeneity than EVs interacting with bEnd.3 cells (P = 0.013), indicating more uniform diffusional behavior following uptake (Figure 3d). In contrast, no significant differences in heterogeneity were observed when data were grouped by EV origin (Figure 3e).

To identify the variables driving separation in the PCA space, we examined the variables that contributed most to PC1 and PC2 (Supplementary Figure 13). EVs interacting with bEnd.3 cells were significantly more time occupied in a confined diffusion state compared with EVs interacting with both ARPE-19 (P = 2.8 × 10^−4^) and HEK293-H cells (P = 1.1 × 10^−6^), suggesting that a larger fraction of EVs reside as membrane-bound particles either at the cell membrane or within intracellular compartments. Conversely, EVs in HEK293-H cells spent significantly more time in the subdiffusive state compared to ARPE-19 (P = 0.0047) and bEnd.3 cells (P = 1.4 × 10^−7^), consistent with EVs being significantly more prone to move within densely crowded cytosolic environments in HEK293-H (Figure 3f, g).

Together, these results point towards the intracellular trafficking behavior of EVs being largely governed by the recipient cell environment rather than the origin of the EVs. EVs interacting and internalized by HEK293-H cells exhibit relatively homogeneous diffusional behavior dominated by cytosolic motion, whereas EVs in bEnd.3 cells display more heterogeneous dynamics with a greater fraction of EVs residing at the cell membrane or within membrane-associated compartments. However, while the fraction of time spent in each diffusional state provides direct insight into intracellular motion at the single vesicle level, it fails to reveal how EVs transitions between these states during trafficking.

### Diffusional transition probabilities further reveal cell-type-dependent EV trafficking behavior

To further characterize EV trafficking behavior, we mapped transitions between diffusional states along individual trajectories at the single-vesicle level to capture time-dependent diffusional fingerprints (Figure 4a and Supplementary Figure 14-16). From each trajectory, we calculated the probability of transitions between the four diffusional states for each EV type in each recipient cell type across multiple time points (Figure 4b and Supplementary Figure 17-19). This analysis captures how EVs switch between distinct diffusional and cellular interacting behavior, for example, transitioning from cytosolic diffusion to confined states associated with membrane-bound compartments, and therefore provides a dynamic dimension of intracellular trafficking pathways.

When grouped by recipient cell type, the transition probabilities showed partial separation in a PCA (Figure 4c), indicating that the temporal progression of diffusional transitions is largely determined by the cellular environment. Analysis of heterogeneity between recipient cell types further supported this observation. The transition probabilities seen from EVs in HEK293-H cells were significantly more homogeneous than those observed in bEnd.3 cells (P = 6.2 × 10^−5^), suggesting that EV trafficking follows more uniform intracellular pathways in HEK293-H cells (Figure 4d). In contrast, EV behavior in bEnd.3 cells displayed greater diversity in motion, indicating a more dynamic transition between different intracellular trafficking routes.

Inspection of the PCA loadings revealed that separation between cell types was primarily driven by transitions involving subdiffusive and confined states. EVs in bEnd.3 cells were significantly more likely to transition from subdiffusive motion, consistent with diffusion in crowded cytosolic environments, to confined motion associated with membrane-bound compartments. In contrast, EVs in HEK293-H and ARPE-19 cells more frequently transitioned from subdiffusive motion to normal diffusion, indicating continued movement within less restricted cytosolic regions (Supplementary Figure 20).

When transition probabilities were grouped by EV origin, no significant differences in heterogeneity or in the principal contributors to PC1 and PC2 were detected, further indicating that EV origin has minimal influence on intracellular trafficking and temporal progression of diffusional transitions (Supplementary Figure 20).

Together, these findings demonstrate that intracellular EV trafficking, deconvoluted at the single-EV level, is largely determined by the recipient cell environment rather than the biological origin of the EVs. EVs internalized by HEK293-H cells exhibit relatively uniform trajectories and progression transition characterized by dynamic cytosolic diffusion and frequent transitions to non-confined states, suggesting a well-defined intracellular transport pattern (Figure 4f). In contrast, EVs in bEnd.3 cells display highly heterogeneous trajectories with frequent transitions to confined states, consistent with multiple trafficking pathways involving membrane-associated compartments. ARPE-19 cells exhibit intermediate behavior, with trajectories that remain predominantly dynamic but display moderate variability between individual EVs.

## Discussion

We systematically examined how extracellular vesicle (EV) origin and recipient-cell identity influence intracellular EV trafficking by combining single-vesicle tracking with single-particle diffusion-state classification. Across EVs derived from four sources and analyzed in three recipient cell types, EV origin had only a modest and non-significant effect on intracellular dynamics after uptake, whereas recipient-cell identity was the significant and principal determinant of EV behavior. These findings argue against a unidirectional model in which intracellular EV fate is specified predominantly by vesicle origin and instead support a framework in which EV trafficking is determined by an interaction between EV-associated properties and the intracellular transport environment of the recipient cell.

These observations have important implications for the interpretation of EV-mediated communication. A mechanistic understanding of how EVs are processed after uptake is essential for defining their roles in physiology and disease, yet our results indicate that these processes cannot be inferred from EV origin alone. Rather, the intracellular context of the recipient cell appears to be a major determinant of how EVs move and, potentially, how their functional effects are realized, which is consistent with previous reports demonstrating no clear clustering based on cell source (15). This has relevance for strategies aimed at functionally modifying or engineering EVs for therapeutic use, as it suggests that recipient-cell-specific trafficking programs may critically shape the outcome of EV delivery (22-24).

Our findings also underscore the utility of single-vesicle approaches for resolving the determinants of intracellular EV behavior. Single-vesicle and trajectory-based analyses have previously revealed heterogeneous motion states and trafficking behaviors for nanoparticles and vesicular structures in living cells (22, 24, 32-34). These studies further report dynamic transitions between distinct motional states during intracellular EV transport, including characteristic “slow-fast-slow” trafficking patterns (22). Consistent with this, EVs on astrocytes showed more confined motion, whereas EVs on microglia displayed more dynamic diffusional patterns (25). Extending these approaches across multiple EV sources and recipient-cell contexts, our study shows that this heterogeneity is explained primarily by the identity of the recipient cell rather than by the source of the EV. Similar observations have previously been reported using endpoint measurements by immunofluorescence microscopy (35), indicating that related trends can be detected at the population level, although such approaches lack the temporal resolution required to resolve dynamic EV behavior. Similarly, three-dimensional holographic fluorescence imaging and related single-vesicle approaches have shown that EVs undergo transitions between diffusive and directed transport in live cells, supporting the view that intracellular trafficking is dynamic and heterogeneous (6). Complementary studies of EV-cell interactions further demonstrate that while binding and fusion kinetics can be similar across recipient cell types, the overall efficiency of uptake and fusion varies substantially, emphasizing the dominant role of the recipient-cell environment in shaping EV behavior (23).

A major advantage of single-particle tracking is that it enables a dynamic insight into individual vesicles, revealing transitions between heterogeneous trafficking behaviors masked in ensemble measurements or low-temporal-resolution imaging (26, 36). However, this technique is limited to tracking detected EVs; consequently, EVs that either remain below the detection threshold, do not interact with the cell, or move too fast to be tracked are not captured in the dataset. Thus, our measurements only quantify the fraction of EVs that interact with or enter the cell.

An important next step will be to investigate possible associations between intracellular behavior and functional readout, such as cargo delivery or interaction with intracellular structures such as actin networks, endosomal compartments, or other organelles (23, 26). Integrating trajectory analysis with molecular markers of intracellular compartments could provide further mechanistic insight into the pathways underlying the diffusion states observed in this study. Such information may also help explain differences in delivery efficiency between cell types. Our observed differences in the intracellular trafficking of HEK293-H and bEnd.3 cells may further explain why the latter are more difficult to transfect. Brain endothelial cells, such as bEnd.3 cells possess specialized barrier properties and complex endo-lysosomal systems, and are known to be a difficult target for non-viral transfection (37, 38). Consistently, EVs in bEnd.3 cells exhibit heterogeneous and frequently confined trajectories, suggesting routing through membrane-bound compartments, where cargo may be degraded. In contrast, EVs in HEK293-H cells display more uniform and dynamic diffusion, indicative of a more permissive intracellular environment that facilitates cytoplasmic access and could explain why HEK293-H cells are easily transfected (39).

Beyond native EV research, quantitative single-vesicle analysis may contribute to the development of EV-based therapeutic strategies. Understanding how different recipient cell types process incoming EVs could guide design-rules for functional EV-based delivery systems with improved targeting and cytoplasmic or nuclear delivery efficiency. More broadly, advances in high-throughput imaging and analysis platforms are likely to enable large-scale screening of EV-cell interactions, facilitating the systematic investigation of EV behavior across diverse experimental conditions.

## Materials and Methods

### Cell Culture

Human retinal pigment epithelial cell line ARPE-19, mouse brain endothelial cell line bEnd.3, and murine myoblast cell line C2C12 were obtained from ATCC (Manassas, VA, USA). The human embryonic kidney cell line was purchased from Thermo Fisher Scientific (Waltham, MA, USA). ARPE-19 cells were cultured in Dulbecco’s modified Eagle’s medium/F-12 (DMEM/F-12; Thermo Fisher Scientific), whereas HEK293-H, bEnd.3 and C2C12 cells were maintained in Dulbecco’s Modified Eagle Medium (DMEM) supplemented with 10% heat-inactivated fetal bovine serum (FBS), 2 mM L-glutamine, and 1% penicillin-streptomycin (PS) (Thermo Fisher Scientific, Waltham, MA, USA). Cultures were incubated at 37 °C in a humidified atmosphere containing 5% CO_2_.

For EV production, cells were grown to approximately 90% confluence in T175 flasks. Prior to EV collection, cultures were washed twice with 10 mL phosphate-buffered saline (PBS) to minimize contamination from serum-derived vesicles. Cells were then incubated in 20 mL EV collection medium (DMEM, supplemented with 10% EV-depleted FBS and 1% PS) per flask for 24 h before harvesting conditioned medium.

### C2C12 EV Isolation

Conditioned medium was sequentially centrifuged to remove cells and debris: 300x g for 10 min, 2,000x g for 20 min, and 15,500x g for 30 min (all at 4 °C). The clarified supernatant was filtered through 0.2 μm membranes to eliminate large particles.

EVs were pelleted by ultracentrifugation at 100,000x g for 2 h at 4 °C using an Optima L-80-XP ultracentrifuge with a 60Ti rotor (Beckman Coulter). Pellets were resuspended in 200 μL PBS.

For further purification, samples were processed using qEV original-35 size-exclusion chromatography (SEC) columns according to the manufacturer’s instructions. Fractions (0.5 mL each) were collected and analyzed for protein concentration using the Qubit Protein Assay Kit (Thermo Fisher Scientific) and for particle size and concentration by nanoparticle tracking analysis (NTA). EV-enriched fractions (typically fractions 1-5 based on particle-to-protein ratio) were pooled, aliquoted (total volume 2.5 mL), and stored at ™80 °C until further use.

### Milk-Derived EV isolation

Fresh bovine milk was processed immediately upon receipt and kept at 4 °C throughout. To remove the cream layer, milk was centrifuged at 1,500x g for 20 min at 4 °C. The upper fat layer was discarded, and the underlying skim milk was carefully recovered.

Skim milk was further clarified by centrifugation at 20,000x g for 1 h at 4 °C to remove residual fat globules, casein aggregates, and debris. The resulting supernatant (milk serum) was transferred to clean ultracentrifugation tubes.

Milk serum was ultracentrifuged at 100,000x g for 2 h at 4 °C. The EV-containing pellet was resuspended in sterile PBS and further purified by SEC using qEV original-35 columns. Fractions (0.5 mL) were collected and analyzed for protein concentration (Qubit assay) and particle size and concentration (NTA). Fraction 2 consistently showed EV enrichment with minimal protein contamination and was therefore collected, aliquoted, and stored at ™80 °C for downstream analyses.

### Plasma-Derived EVs Isolation

#### Plasma Collection

Blood sampling was performed as previously described (40). Briefly, an 18G Venflon (BD Vialon, Becton Dickinson, NJ, USA) was inserted into a cubital vein 30 minutes prior to collection of the first blood sample. Blood was collected in BD Vacutainer K2 EDTA tubes (Becton Dickinson). To obtain cell- and platelet-free plasma, samples were centrifuged at 1500x g for 15 minutes at 20 °C in a swing-bucket centrifuge without brake. The plasma supernatant was carefully transferred to a new tube without disturbing the buffy coat and centrifuged again under the same conditions. Subsequently, the supernatant was cooled on ice and centrifuged at 16,000x g for 10 minutes at 4 °C to remove residual platelets and cellular debris, and then stored at -80 °C. Hemolysis was assessed by measuring free hemoglobin at an absorbance of 414 nm, and samples with A414 > 0.2 were excluded to avoid contamination of extracellular vesicle preparations by lysed cells.

Retro-orbital venous blood from mice was collected in BD Vacutainer K2 EDTA tubes (Becton Dickinson). Plasma was separated by centrifugation at 1,900x g for 10 min at 5 °C, aliquoted, and stored at ™80 °C until analysis.

#### EV Isolation

Prior to EV isolation, plasma samples were thawed at 4 °C and centrifuged at 15,000x g for 15 min at 4 °C to remove residual cellular debris.

EVs were isolated from 500 μL of cleared plasma using size-exclusion chromatography (qEV columns, Izon Bioscience, New Zealand) according to the manufacturer’s protocol. Fractions (0.5 mL each) were collected and evaluated for protein concentration using the Qubit Protein Assay Kit and for particle size and concentration by NTA.

EV-enriched fractions with minimal protein contamination were pooled and concentrated using 10 kDa molecular weight cut-off ultrafiltration devices (Amicon Ultra-2, Merck Millipore, Germany) to a final volume of 100 μL. Concentrated EV preparations were subsequently characterized by NTA.

### Western Blotting of EVs

Cells were lysed in 400 μL RSB300-U buffer (300 mM NaCl, 2.5 mM MgCl_2_, 10 mM Tris, pH 7.4, 1 M urea, 0.5% Triton, and 40 μg mL^−1^ protease inhibitor). For EV western blotting, protease inhibitor was added after EV purification. Protein concentrations of cell lysates and EVs were measured using the Qubit Protein Assay Kit (Thermo Fisher Scientific), and 30 μg protein was loaded per sample. Samples were prepared in 4x NuPAGE LDS sample buffer (Invitrogen, Waltham, MA, USA) and were resolved on 4-12% NuPAGE Novex Bis-Tris gels (Invitrogen) with a 10-250 kDa protein ladder, run at 200 V for 35 min, and transferred to PVDF membranes at 30 mA for 1 h at 4 °C. Membranes were probed with antibodies against CD81 (sc-166029, Santa Cruz Biotechnology; diluted 1:100),), Calnexin (ab22595, Abcam; diluted 1:500), flotillin 1 (1:500, 610821, BD Biosciences, CA, USA), blocked in 5% milk, and incubated with HRP-conjugated secondary antibody (1:5,000; DAKO, Agilent). Signals were detected using Pierce ECL Plus, recorded on X-ray film, and developed with an Agfa CP1000 developer.

### Nanoparticle tracking analysis (NTA) of EVs

Particle size and concentration were determined by nanoparticle tracking analysis (NTA) using a NanoSight LM10 (Malvern Instruments Ltd., Malvern, UK). Samples were diluted in PBS prior to analysis and measured in triplicate each for 60s at RT with a camera level of 16, and a detection threshold of 5. Videos were recorded and analyzed using the NTA software (version 3.1 build 3.1.45).

### Transmission Electron Microscopy (TEM) Imaging

EVs were imaged by transmission electron microscopy using an FEI Tecnai G2 Spirit equipped with a bottom-mounted TVIPS CMOS 4k camera (TEM-cam-F416). Carbon film-coated 400-mesh copper grids were glow discharged for 45 s, incubated twice with 3 μL EV suspension for 1 min each, and blotted dry with Whatman qualitative filter paper after each incubation. Grids were then stained with 3 μL of 2% uranyl formate for 5 s and blotted dry.

### Cy5 loading of EVs using electroporation

All EVs were treated at a concentration of 2 × 10^10^ EVs in a total for electroporation accommodated in trehalose pulse medium (50nM in PBS) (Cat. No. T0167) and between 20 to 200 pmol siRNA-Cy5. The final volume is 40 μL with a final Trehalose concentration of 50mM. Electroporation was performed on a GenePulser II Electroporation System with capacitance extender (Bio-Rad, Hercules, CA, USA) using 0.4 cm cuvettes with aluminum electrodes (Cat. No. 165-2086). Cool the cuvette before inserting and mix the EV with siRNA and put on ice for 1min. All EVs were treated with either 400V 1x to 5x pulses and 750V 1x to 5x. After electroporation, the mixture was incubated at 37 °C for 30 min and then at 4 °C for overnight(30).

### Cellular uptake studies and imaging acquisition

EV concentrations were quantified by NTA to ensure comparable particle numbers across conditions before electroporation. Recipient cells were seeded in 8-well μ-slide (ibiTreat ibidi) at densities of 8 × 10^4^, 1 × 10^5,^ and 5 × 10^4^ per well in 300 μL culture medium for HEK293-H, ARPE-19, and bEnd.3 cells, respectively, followed by overnight incubation. Cells were then treated with 4 × 10^10^ EVs (40 μL) per well from each source loaded with Cy5-siRNA through electroporation. For control experiment Cy5-siRNA only samples were prepared by electroporation in the absence of EVs. The cell nuclei were stained with Hoechst 33342 (Molecular Probes) for 20 min at 37 °C, followed by fixation with 4% PFA for 10 min and washed with PBS. The plasma membrane was stained with wheat germ agglutinin-Alexa Fluor 488 (Invitrogen, W11261) for 10 min at room temperature. Cells were then washed twice with PBS before being visualized by confocal laser scanning microscopy (LSM 710, Zeiss, Germany) and analyzed using ZEISS Zen Software.

### Acquisition of EV loading TIRF imaging data

All single-EV experiments were performed on an Oxford Nanoimager (Oxford, UK) operated in TIRF mode. Biotinylated, membrane-labelled EVs were immobilized in microscope surface chambers prepared on clean glass slides by oxygen plasma and ultraviolet ozone treatment for 1 h. Surfaces were then functionalized for 30 min with PLL-g-PEG and PLL-g-PEG-biotin (100:1), washed with Milli-Q water, and assembled with a sticky-Slide VI 0.4 chamber (ibidi), as described previously. A neutravidin layer (0.1 g L^−1^) was added for 30 min before imaging, followed by washing with PBS (30, 41).

EVs were electroporated as described previously. The electroporation was followed by EV membrane labelling using 300 ATTO 488-DOPE lipids and 300 DSPE-PEG(2000)-Biotin lipids per EV, following previously published protocols (30, 42). EVs were purified from free labels using 100kDa Amicon Spin filters for 3x rounds at 3.4 × 1000 g with a total flow-through volume of 1.5 mL. The labelled and electroporated EVs were introduced into the chamber and allowed to immobilize for 5 min, yielding ∼100-200 vesicles per field of view. Unbound EVs were removed by exchanging the chamber volume 5 times with PBS using a peristaltic pump. Imaging was performed using 100 ms exposure time and alternating laser excitation using 1% 488 nm laser and 15% 640 nm laser.

Membrane fluorescence and the corresponding signal from lumen-encapsulated siRNA were colocalized, integrated, and background-corrected for individual EVs in both imaging channels across all fields of view using in-house Python software for localization and tracking. This enabled automated analysis with nanometre-scale colocalization precision (43).

### Acquisition of real-time imaging data

Recipient cells were seeded in μ-Slide VI ibidiTreat imaging channels at densities of 1 × 10^5^, 1 × 10^5,^ and 3 × 10^5^ per well in 100 μL culture medium for HEK293-H, ARPE-19, and bEnd.3 cells, respectively, followed by overnight incubation. Live tracking experiments of extracellular vesicles were conducted using an Oxford Nanoimager microscope (Oxford, UK), as described earlier(36). In short: Imaging was performed with a 100x oil-immersion objective (NA 1.4) at 37 °C. Samples were delivered using an external peristaltic pumping system at a flow rate of 0.1 mL min^−1^. Dual-channel images were acquired simultaneously at 16-bit grayscale with a frame size of 428 × 684 pixels per channel, corresponding to an effective field of view of approximately 50 μm × 80 μm. All measurements were carried out under highly inclined laminated optical sheet (HILO) illumination.

### Data analysis

#### Particle tracking

Single-particle trajectories were obtained using an in-house developed Python software (36). Intensity-based outliers were excluded using a robust modified z-score criterion computed from the median and median absolute deviation. Trajectories were subsequently segmented and classified using DeepSPT (26). Fractional diffusion states and transition probabilities were calculated using Python. See SI for data and code availability.

### Multivariate and statistical analysis

Fractional diffusion features and transition probabilities were analyzed in R. Principal component analysis (PCA) was performed using the *mdatools* package, and scores and loadings were visualized with 95% confidence ellipses. Euclidean distances of PCA scores were used for PERMANOVA (999 permutations) and dispersion testing. Group differences in fractional diffusion states and transition probabilities were assessed by one-way ANOVA with Tukey post hoc tests. The P-values for all statistical tests in the main text and the Supporting Information are *P ≤ 0.05, **P ≤ 0.01, and ***P ≤ 0.001. See SI for data and code availability.

### Ethics

All animal experiments were carried out in compliance with the European Directive 2010/63/EU on the protection of animals used for scientific purposes. The study protocol received approval from the Danish Animal Experiments Inspectorate (Dyreforsøgstilsynet) under license number 2022-15-0201-01320. All procedures were conducted at the animal research facilities of Aarhus University (Skou Building). The experiment are performed under Ethical Committee for Region Midtjylland (ref. No. 1-10-72-218-16) and registered in the database clinicaltrials.gov (NCT03380663).

## Acknowledgments

This work was funded by Lundbeck Foundation grant No. R380-2021-1393 for M.G.M. Danish National Research Foundation grant no. 135 (CellPAT) for B.H.B., E.A.S., and J.K. Novo Nordisk Foundation grant NNF23OC0081177 (RNA-META) for H.W., and P.S.

## Author Contributions

M.G.M, Y.Y., and J.K. designed research; M.G.M., Y.Y., and P.S. performed experiments; B.H.B., M.G.M., H.W., and E.A.S. analyzed data; and B.H.B. wrote the paper with inputs from all authors. M.G.M. and J.K. supervised the project.

