## Supplementary Information for "Recipient cell identity governs intracellular transport and membrane interactions of extracellular vesicles across species"

**Figures**

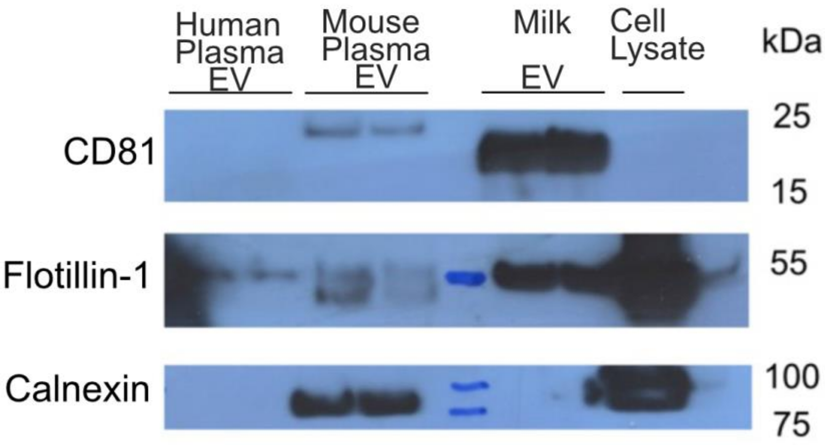

***SI Fig. 1: Western blot of EVs***. *EV markers CD81, Flotillin-1, and Calnexin, from human plasma, mouse plasma, whole cow milk, and cell lysate from C2C12 cells. Western blot of C2C12 is available in previous publication(1).*

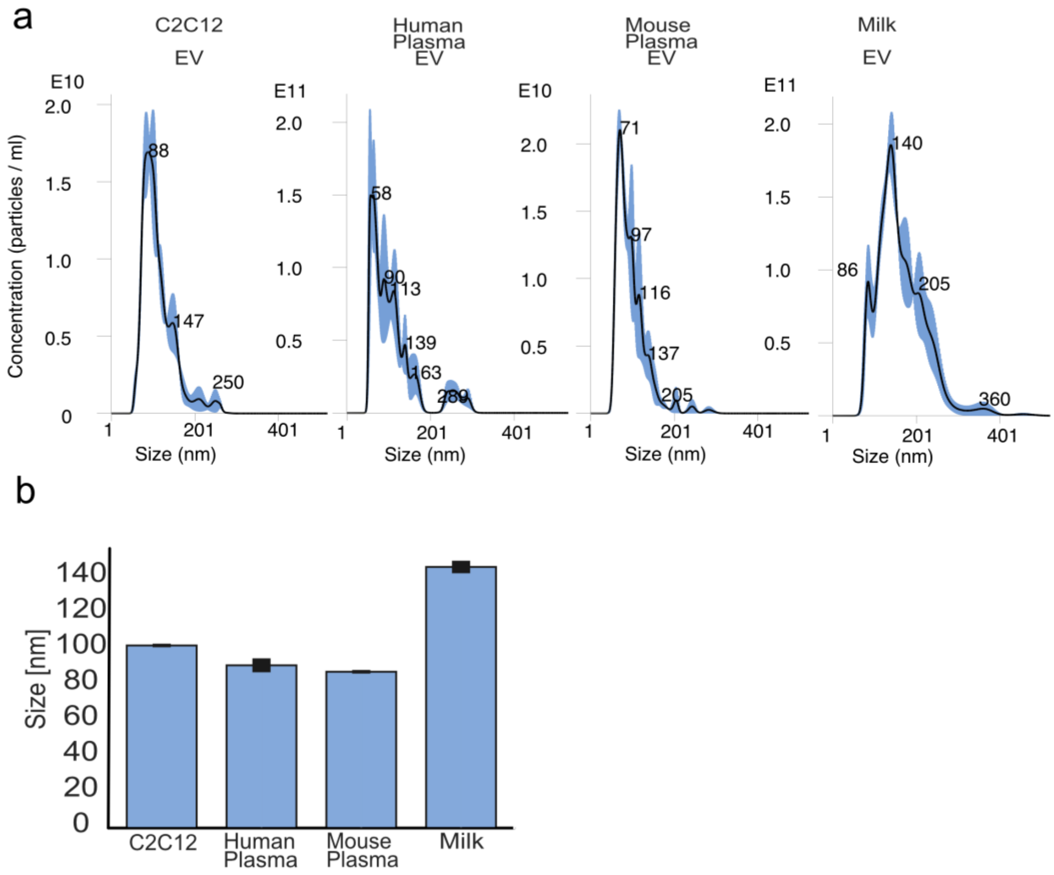

***SI Fig. 2: NTA data of EVs.*** *a) Representative graphs showing the underlying size distribution from NTA measurement. b) Average vesicle size of isolated EVs measured by NTA.*

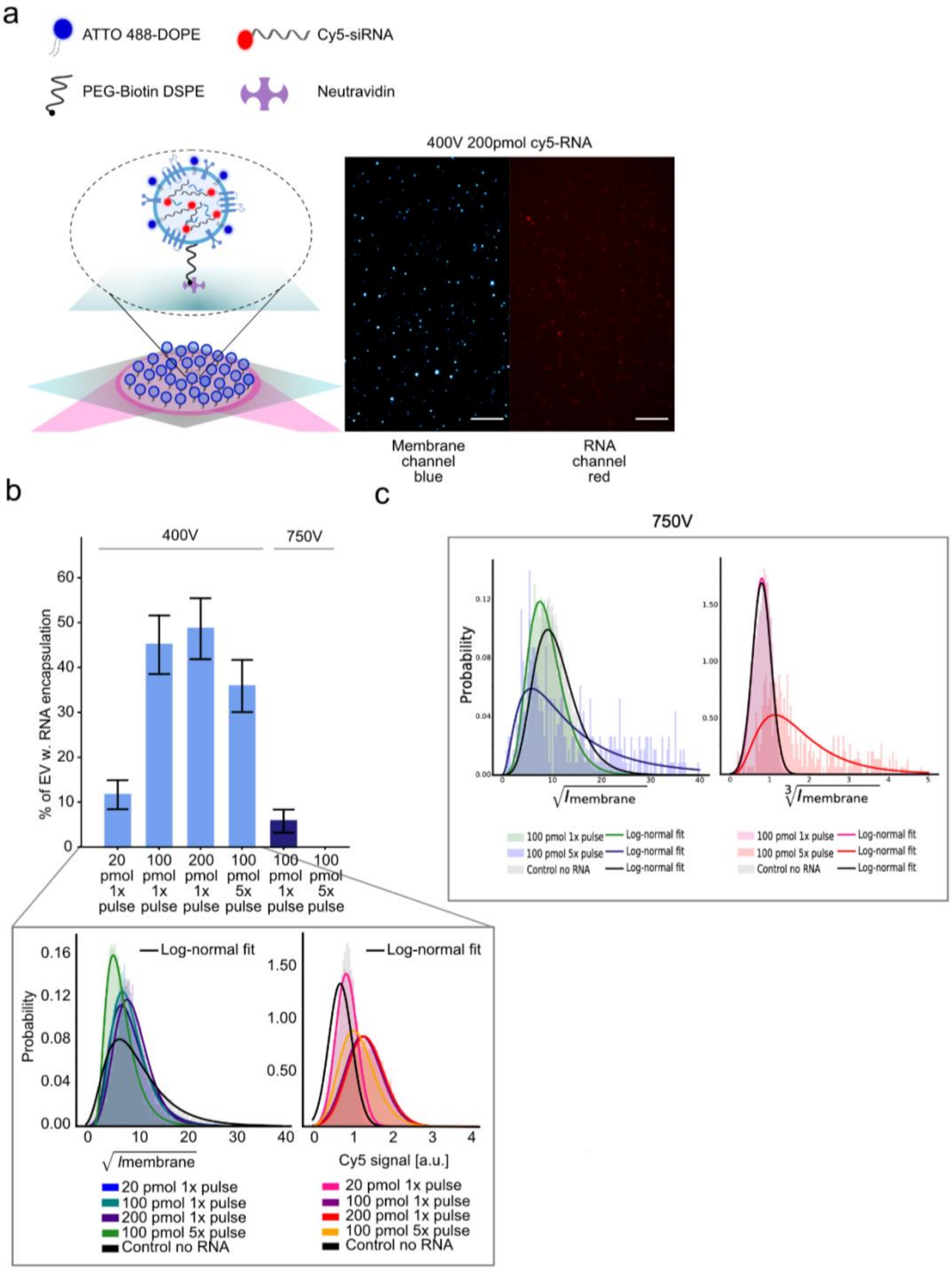

***SI Fig. 3: Single-vesicle quantification and optimized siRNA loading for EV-mediated RNA delivery.*** *a) Single-particle assay to quantify encapsulation efficiency by imaging immobilized EVs membrane-labeled with ATTO 488-DOPE, allowing colocalization of EVs and encapsulated Cy5-siRNA. Representative micrographs showing labeled EV membrane (left) and encapsulated Cy5-siRNA (right). Scalebar 10 µm. b) Quantification of single-vesicle encapsulation efficiency of Cy5-siRNA given in percentage of vesicles, using different loading methods. Zoom-in is showing the underlying membrane signal (left) and Cy5 lumen signal (right). c) Membrane signal (left) and Cy5 lumen signal (right) after electroporation with 750V.*

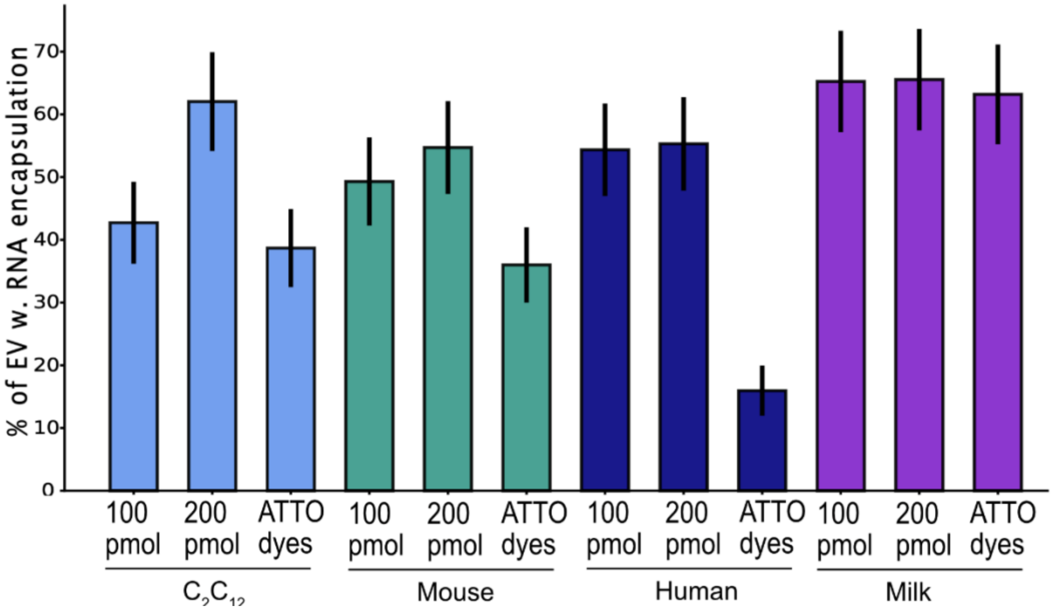

***SI Fig. 4: EV encapsulation efficiency.*** *Quantification of single-vesicle encapsulation efficiency of Cy5-siRNA (or ATTO655-carboxy) given in percentage of vesicles, using different loading methods, in EVs from C2C12 cells, mouse plasma, human plasma, and whole cow milk*.

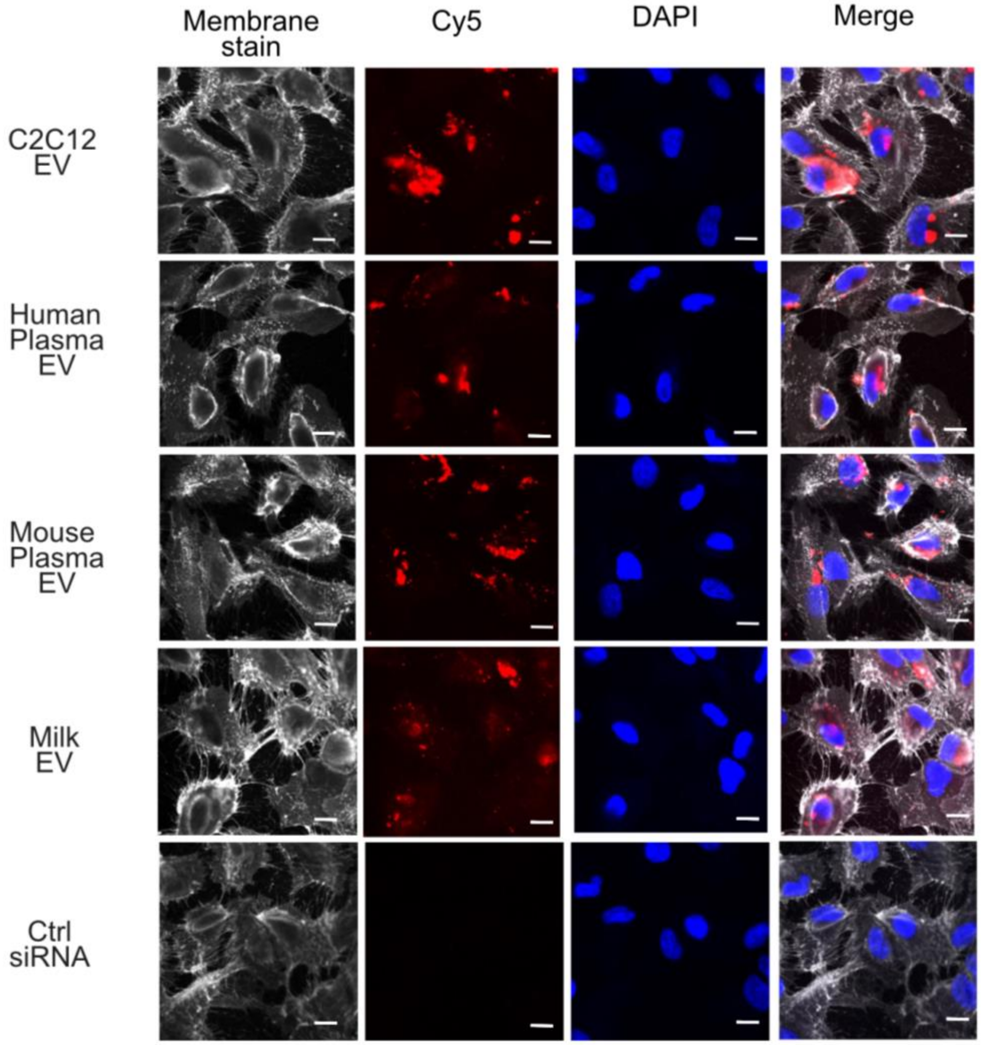

***SI Fig. 5: Cellular uptake of EV delivered Cy5-siRNA in ARPE-19 cells**. Images after 4h, showing representative delivery. Scalebar 10µm.*

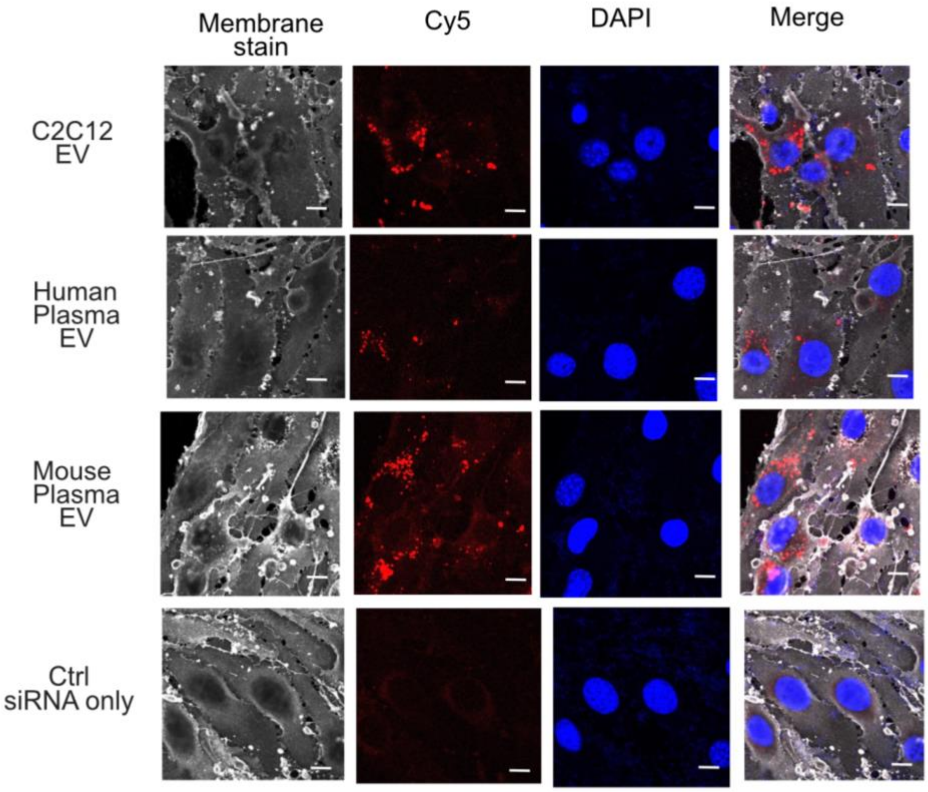

***SI Fig. 6: Cellular uptake of EV delivered Cy5-siRNA in bEnd.3 cells.*** *Images after 24h, showing representative delivery. Scalebar 10µm.*

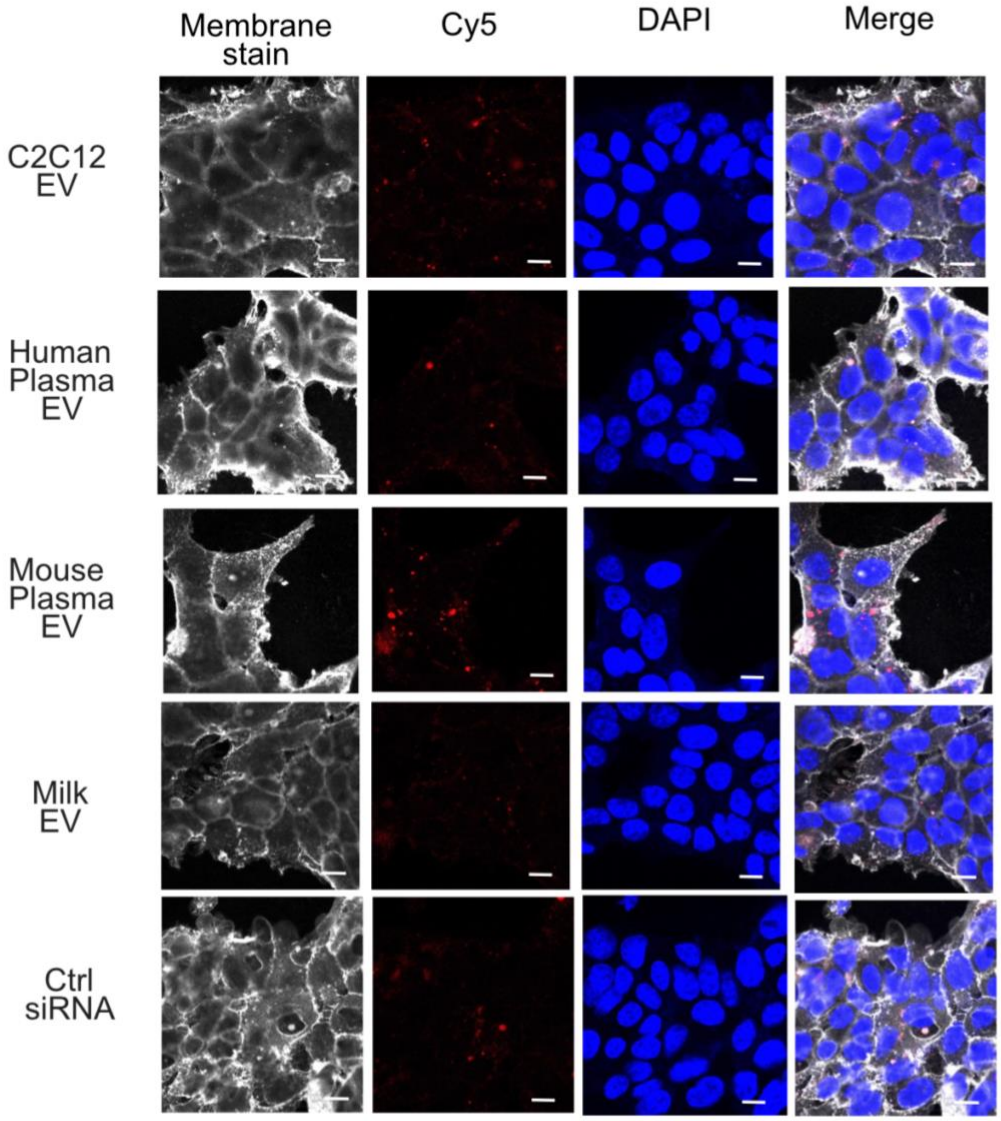

***SI Fig. 7: Cellular uptake of EV delivered Cy5-siRNA in HEK293-H cells.*** *Images after 24h, showing representative delivery. Scalebar 10µm.*

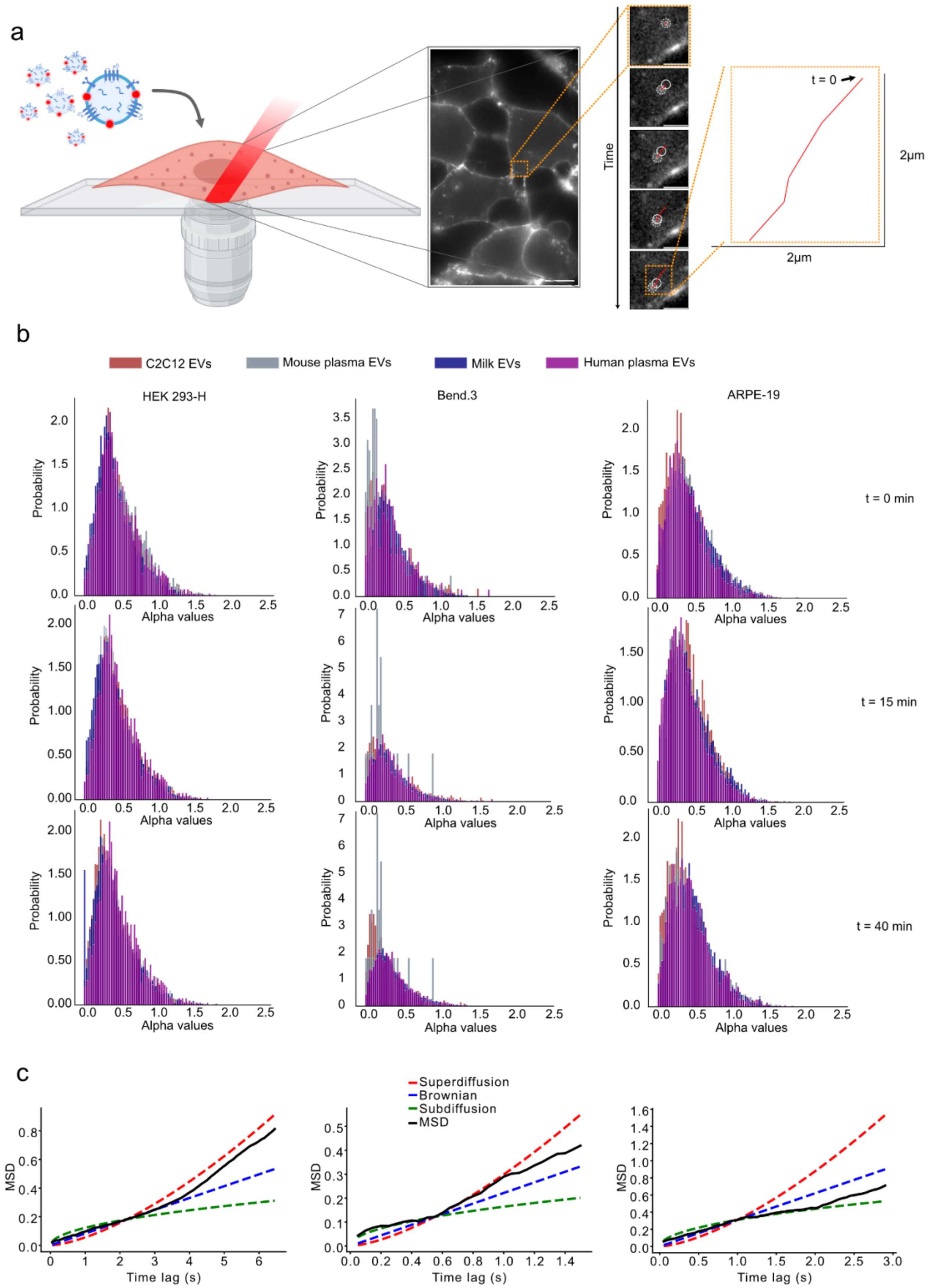

a
Time
t = 0
2μm
2μm
b
C2C12 EVs
Mouse plasma EVs
Milk EVs
Human plasma EVs
HEK 293-H
Bend.3
ARPE-19
Probability
Alpha values
t = 0 min
t = 15 min
t = 40 min
c
Superdiffusion
Brownian
Subdiffusion
MSD
MSD
Time lag (s)

***SI Fig. 8: Single-vesicle tracking for a deconvoluted understanding of uptake.*** *a) Schematic illustration of the experimental setup. The EVs were labeled with ATTO 655-DOPE, enabling real-time tracking of EV movement by obtaining single vesicle trajectories used for quantification of uptake, and diffusional behavior inside cells using HILO microscopy. Scalebar (10$\mu$m), zoom-in (2$\mu$m). b) Comparison of EV behavior between the four different EV types, in different cell types (columns) and different timepoints (rows), to get an insight into the different mechanistic behavior of EVs. This is done by comparing the anomalous diffusion exponent (α) obtained from fitting the trajectory's MSD vs time using $MSD = 4Dt^{\alpha}$, where α ≈ 1 indicates normal diffusion, α < 1 subdiffusion, and α > 1 superdiffusion. c) Representatives of MSD from 3 different EV trajectories.*

a

b

c

d

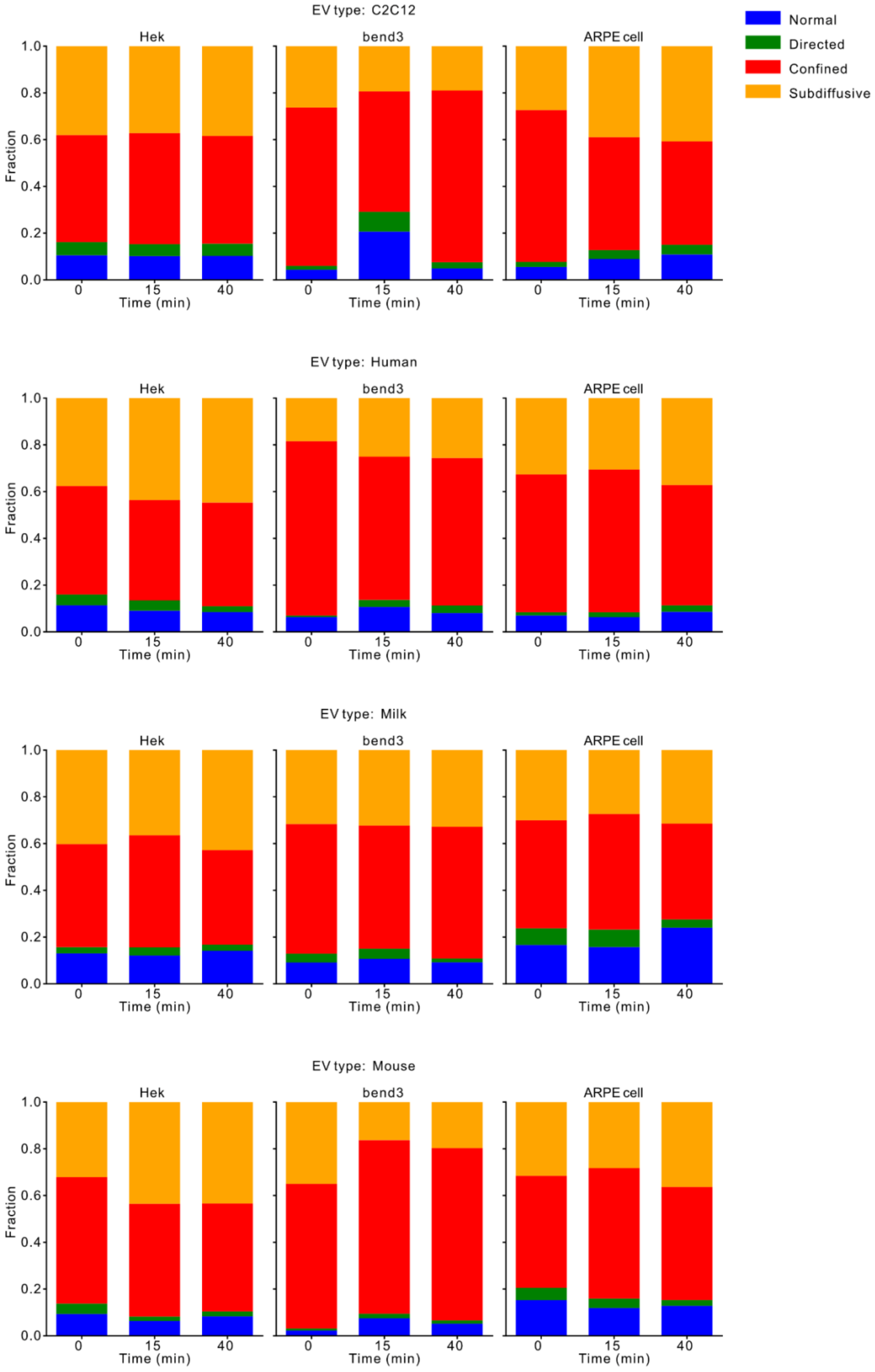

EV type: C2C12
EV type: Human
EV type: Milk
EV type: Mouse
Hek
bend3
ARPE cell
Fraction
Time (min)
0
15
40
0.0
0.2
0.4
0.6
0.8
1.0
Normal
Directed
Confined
Subdiffusive

***SI Fig. 9: Fraction of time spent in each diffusional state.*** *Normal diffusion (blue), directed motion (green), confined motion (red), and subdiffusive motion (yellow). Behavior of EVs from C2C12 cells (a), human plasma (b), whole cow milk (c), and mouse plasma (d) in HEK293-H, bEnd.3, and ARPE-19 cells, measured for 15 minutes at 0, 15, and 40 minutes after addition of EVs to each cell type.*

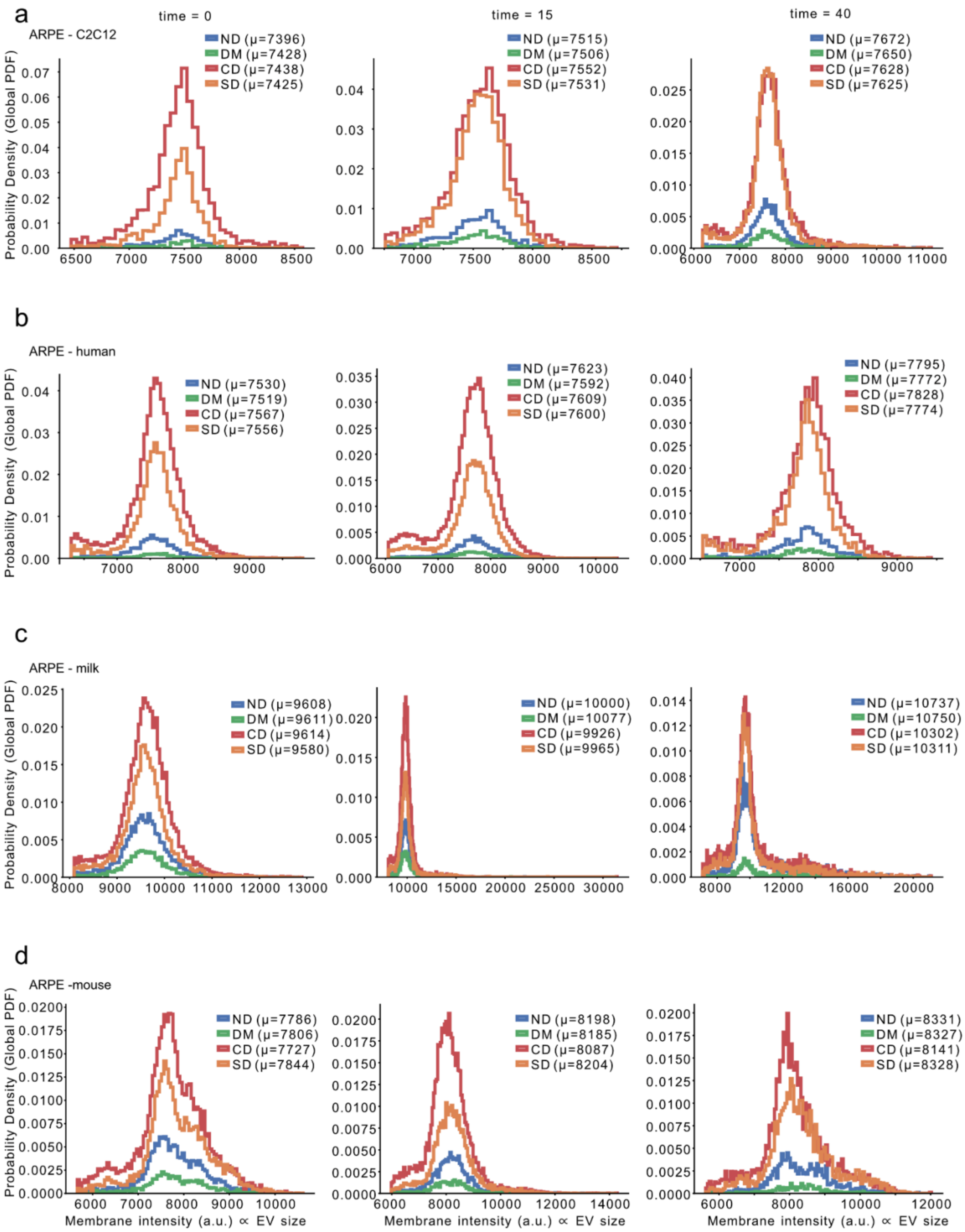
a
time = 0
time = 15
time = 40
ARPE - C2C12
ND (μ=7396)
DM (μ=7428)
CD (μ=7438)
SD (μ=7425)
ND (μ=7515)
DM (μ=7506)
CD (μ=7552)
SD (μ=7531)
ND (μ=7672)
DM (μ=7650)
CD (μ=7628)
SD (μ=7625)
b
ARPE - human
ND (μ=7530)
DM (μ=7519)
CD (μ=7567)
SD (μ=7556)
ND (μ=7623)
DM (μ=7592)
CD (μ=7609)
SD (μ=7600)
ND (μ=7795)
DM (μ=7772)
CD (μ=7828)
SD (μ=7774)
c
ARPE - milk
ND (μ=9608)
DM (μ=9611)
CD (μ=9614)
SD (μ=9580)
ND (μ=10000)
DM (μ=10077)
CD (μ=9926)
SD (μ=9965)
ND (μ=10737)
DM (μ=10750)
CD (μ=10302)
SD (μ=10311)
d
ARPE -mouse
ND (μ=7786)
DM (μ=7806)
CD (μ=7727)
SD (μ=7844)
ND (μ=8198)
DM (μ=8185)
CD (μ=8087)
SD (μ=8204)
ND (μ=8331)
DM (μ=8327)
CD (μ=8141)
SD (μ=8328)
Probability Density (Global PDF)
Membrane intensity (a.u.) ∝ EV size

***SI Fig. 10: Size Distribution of Extracellular Vesicles Across Diffusion States in ARPE-19 cells****. Extracellular vesicles from C2C12 cells (a), human plasma (b), whole cow milk (c), and mouse plasma (d), measured for 15 minutes at 0 (left), 15 (middle), and 40 (right) minutes after addition of EVs to each cell type.*

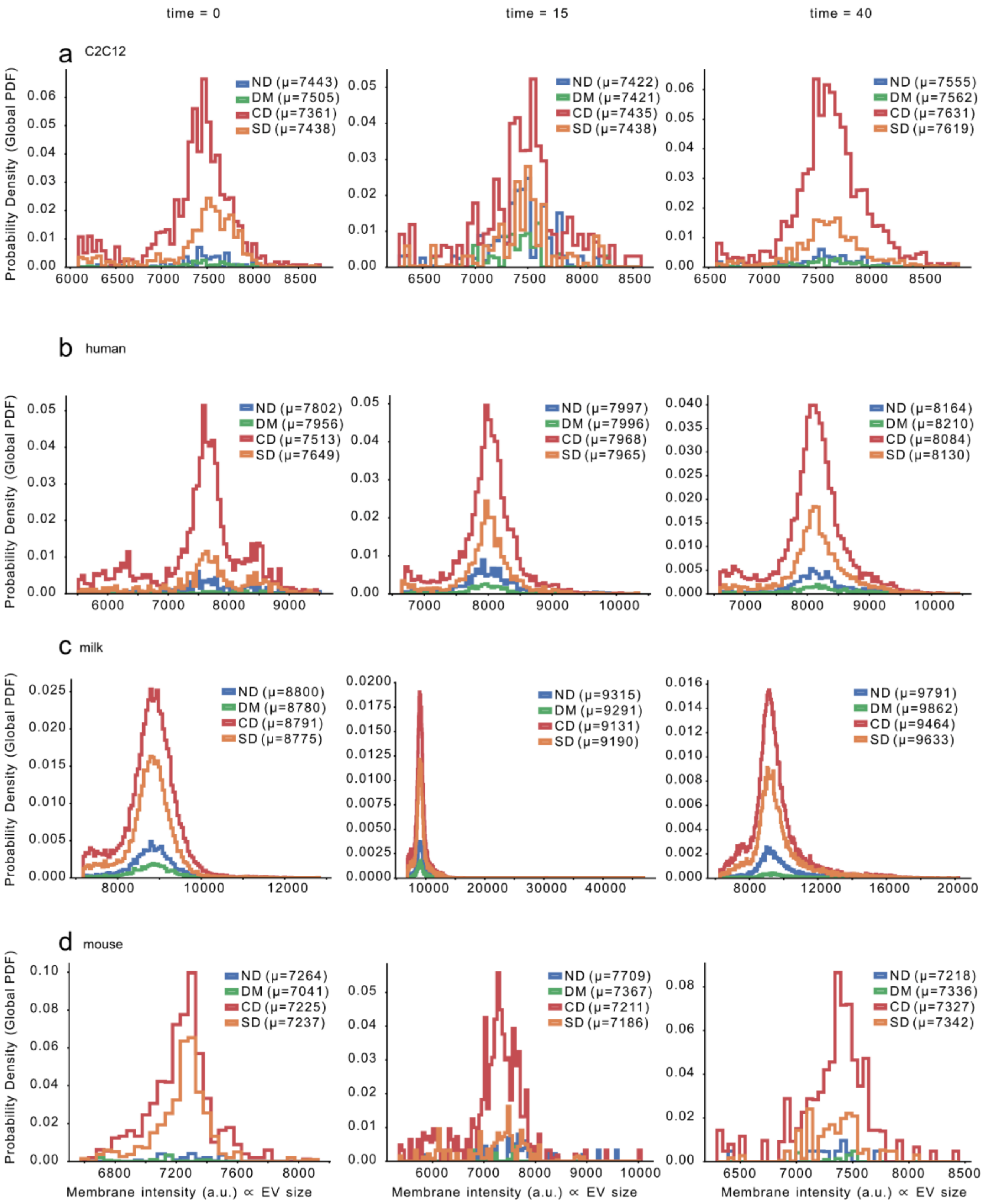
time = 0
time = 15
time = 40
a
C2C12
b
human
c
milk
d
mouse
Probability Density (Global PDF)
ND (μ=7443)
DM (μ=7505)
CD (μ=7361)
SD (μ=7438)
ND (μ=7422)
DM (μ=7421)
CD (μ=7435)
SD (μ=7438)
ND (μ=7555)
DM (μ=7562)
CD (μ=7631)
SD (μ=7619)
ND (μ=7802)
DM (μ=7956)
CD (μ=7513)
SD (μ=7649)
ND (μ=7997)
DM (μ=7996)
CD (μ=7968)
SD (μ=7965)
ND (μ=8164)
DM (μ=8210)
CD (μ=8084)
SD (μ=8130)
ND (μ=8800)
DM (μ=8780)
CD (μ=8791)
SD (μ=8775)
ND (μ=9315)
DM (μ=9291)
CD (μ=9131)
SD (μ=9190)
ND (μ=9791)
DM (μ=9862)
CD (μ=9464)
SD (μ=9633)
ND (μ=7264)
DM (μ=7041)
CD (μ=7225)
SD (μ=7237)
ND (μ=7709)
DM (μ=7367)
CD (μ=7211)
SD (μ=7186)
ND (μ=7218)
DM (μ=7336)
CD (μ=7327)
SD (μ=7342)
Membrane intensity (a.u.) ∝ EV size

***SI Fig. 11: Size Distribution of Extracellular Vesicles Across Diffusion States in bEnd.3 cells.*** *Extracellular vesicles from C2C12 cells (a), human plasma (b), whole cow milk (c), and mouse plasma (d), measured for 15 minutes at 0 (left), 15 (middle), and 40 (right) minutes after addition of EVs to each cell type.*

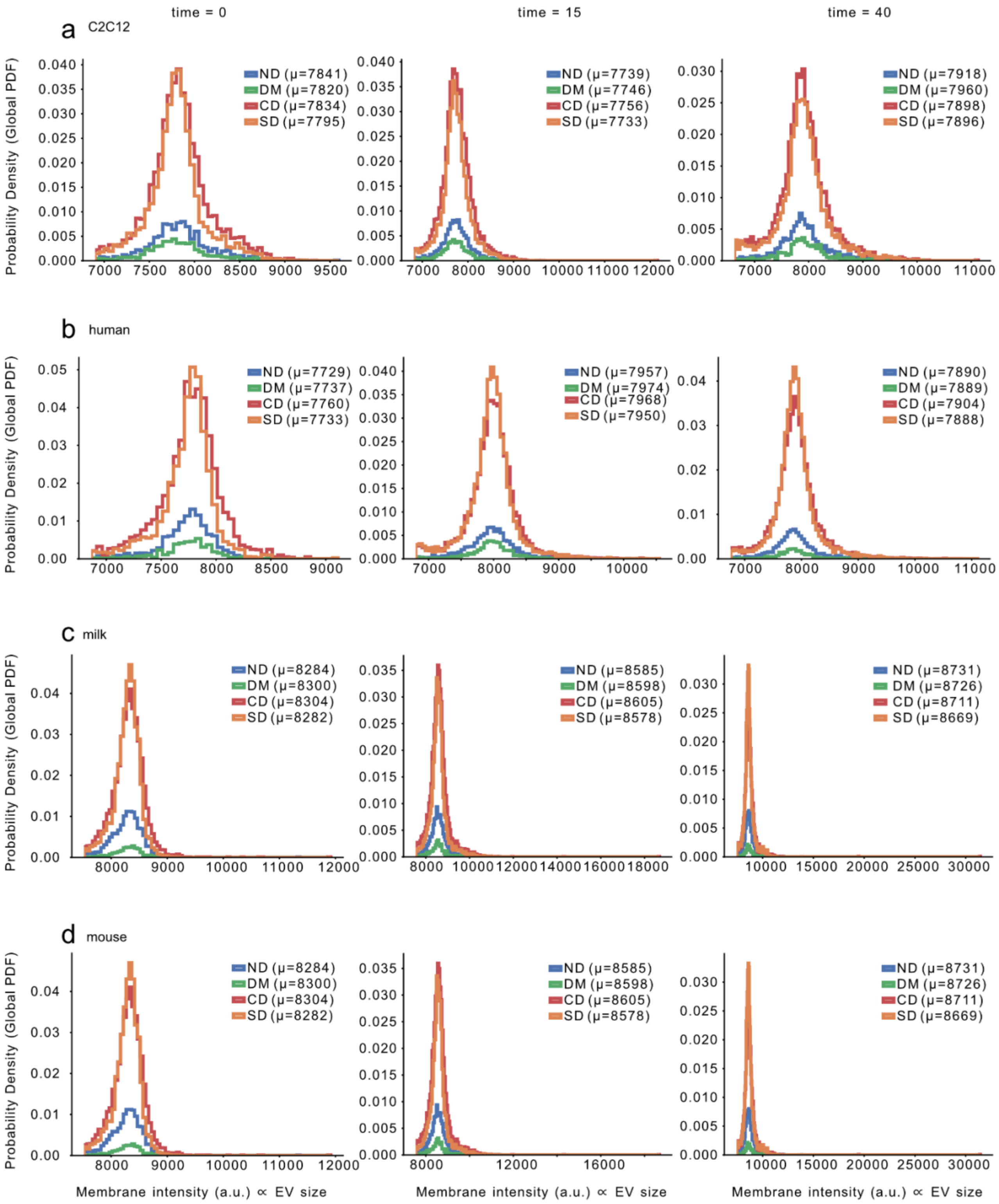
time = 0
time = 15
time = 40
a C2C12
ND (μ=7841)
DM (μ=7820)
CD (μ=7834)
SD (μ=7795)
ND (μ=7739)
DM (μ=7746)
CD (μ=7756)
SD (μ=7733)
ND (μ=7918)
DM (μ=7960)
CD (μ=7898)
SD (μ=7896)
b human
ND (μ=7729)
DM (μ=7737)
CD (μ=7760)
SD (μ=7733)
ND (μ=7957)
DM (μ=7974)
CD (μ=7968)
SD (μ=7950)
ND (μ=7890)
DM (μ=7889)
CD (μ=7904)
SD (μ=7888)
c milk
ND (μ=8284)
DM (μ=8300)
CD (μ=8304)
SD (μ=8282)
ND (μ=8585)
DM (μ=8598)
CD (μ=8605)
SD (μ=8578)
ND (μ=8731)
DM (μ=8726)
CD (μ=8711)
SD (μ=8669)
d mouse
ND (μ=8284)
DM (μ=8300)
CD (μ=8304)
SD (μ=8282)
ND (μ=8585)
DM (μ=8598)
CD (μ=8605)
SD (μ=8578)
ND (μ=8731)
DM (μ=8726)
CD (μ=8711)
SD (μ=8669)
Probability Density (Global PDF)
Membrane intensity (a.u.) ∝ EV size

***SI Fig. 12: Size Distribution of Extracellular Vesicles Across Diffusion States in HEK293-H cells.*** *Extracellular vesicles from C2C12 cells (a), human plasma (b), whole cow milk (c), and mouse plasma (d), measured for 15 minutes at 0 (left), 15 (middle), and 40 (right) minutes after addition of EVs to each cell type.*

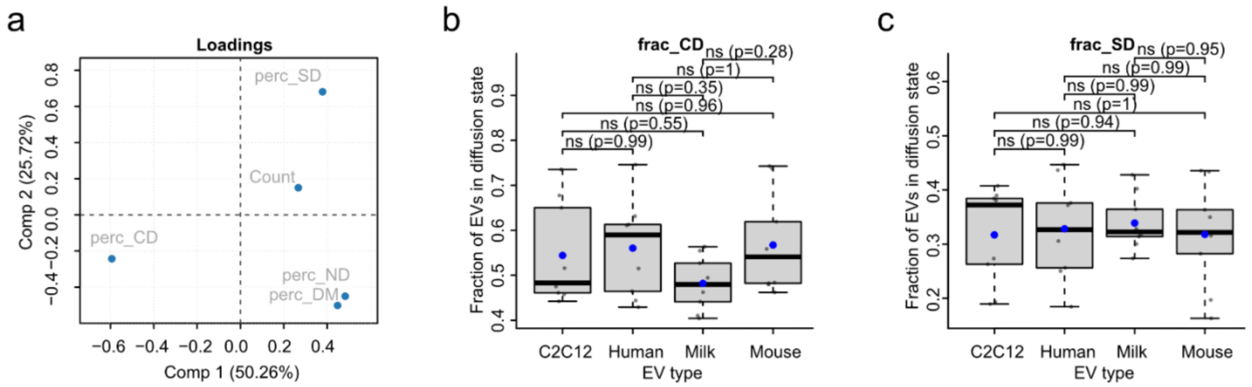

***SI Fig. 13: PCA analysis from fraction of time spent in each diffusional state.*** *a) Loadings describing the driving factors, in PC1 and PC2. b) Boxplot of fraction of EVs in confined state grouped by EV origin. c) Boxplot of fraction of EVs in subdiffusive state grouped by EV origin. Asterisks denote statistical significance (*P ≤ 0.05, **P ≤ 0.01, ***P ≤ 0.001; ns, not significant, P ≥ 0.05), determined by one-way ANOVA with Tukey's post hoc test.*

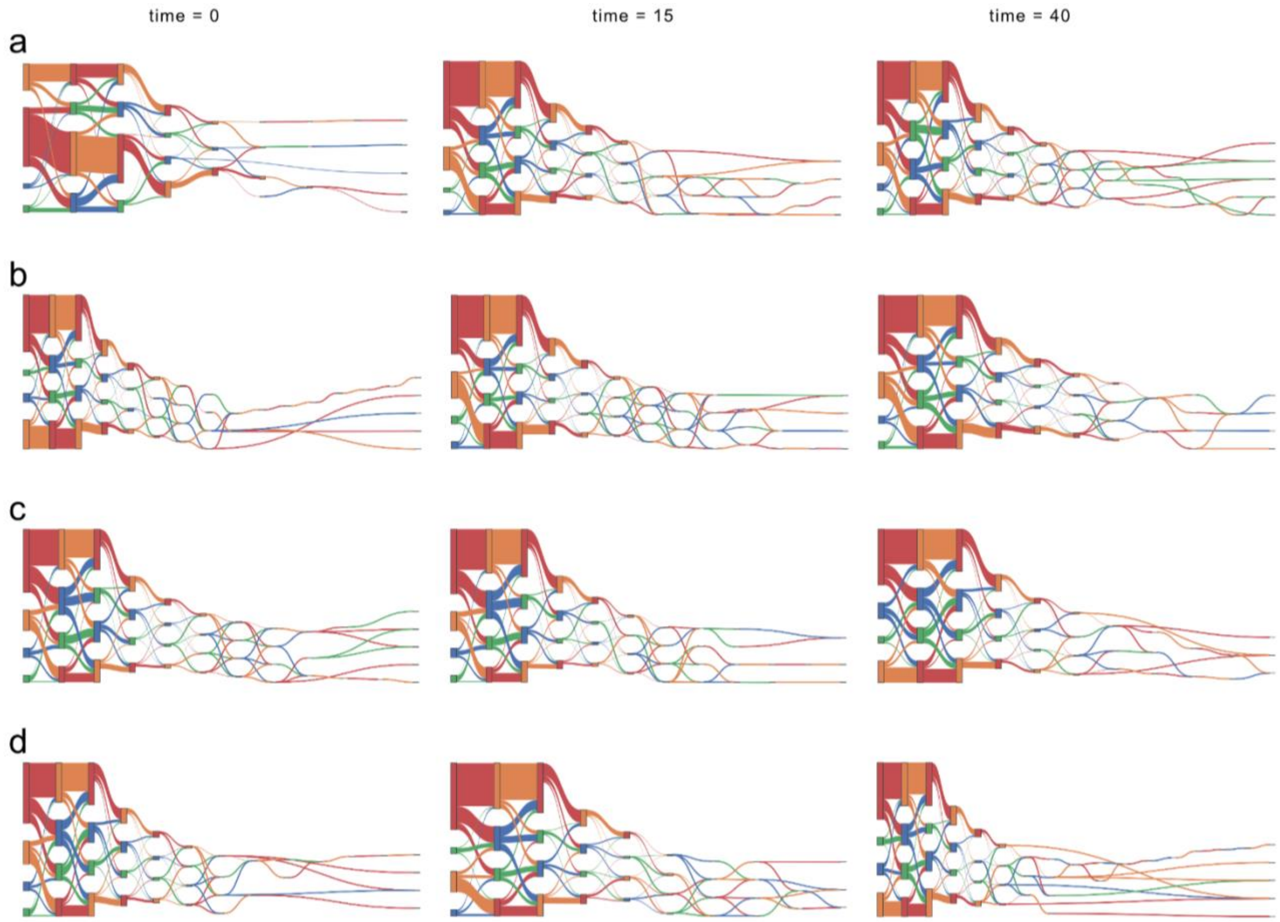

***SI Fig. 14: Sankey diagram of single-particle diffusive behavior over time***. *Graph illustrating transitions between distinct diffusion modes along individual trajectories in ARPE-19 cells. Extracellular vesicles from C2C12 cells (a), human plasma (b), whole cow milk (c), and mouse plasma (d), measured for 15 minutes at 0 (left), 15 (middle), and 40 (right) minutes after addition of EVs to each cell type.*

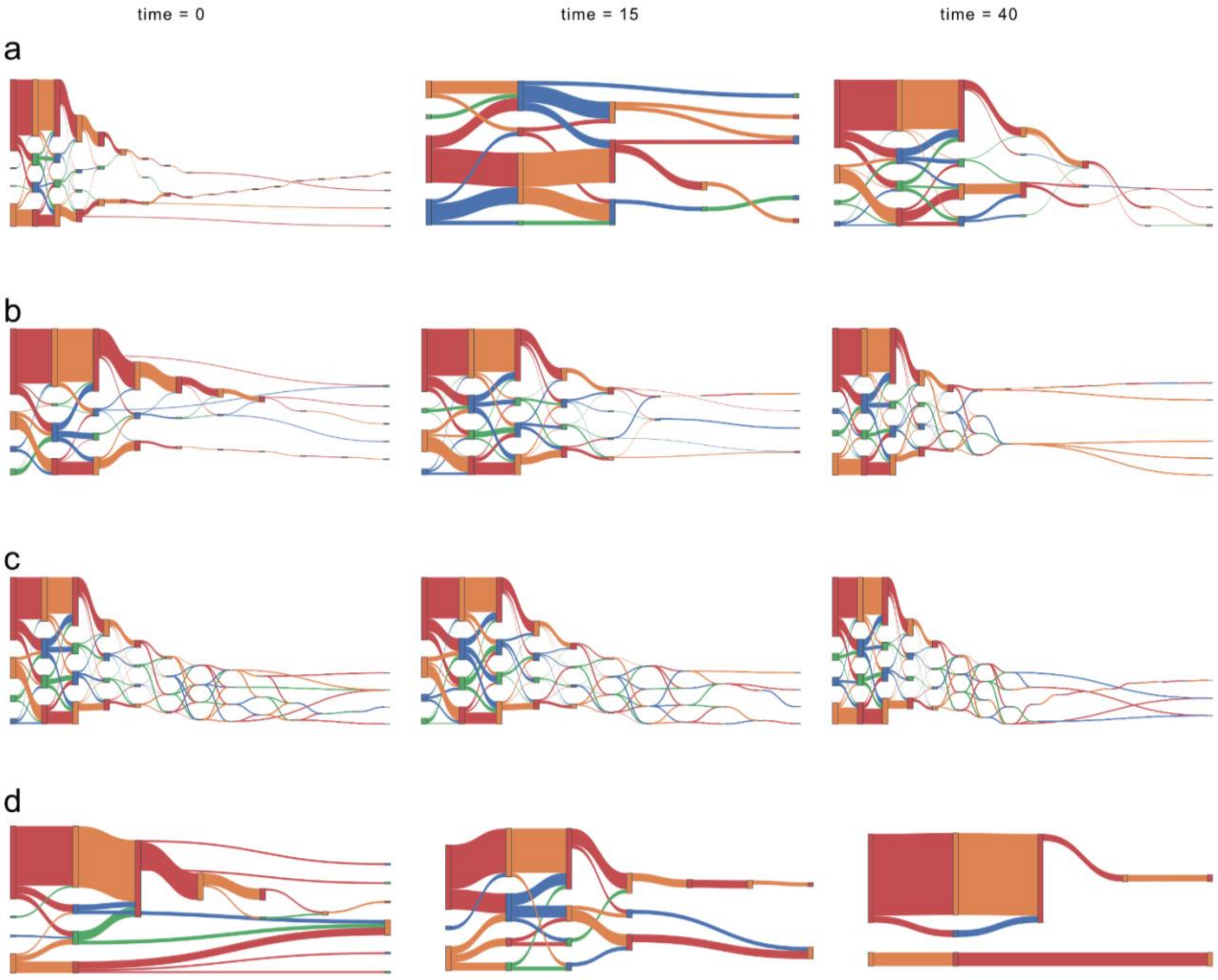

***SI Fig. 15: Sankey diagram of single-particle diffusive behavior over time***. *Graph illustrating transitions between distinct diffusion modes along individual trajectories in bENd.3 cells. Extracellular vesicles from C2C12 cells (a), human plasma (b), whole cow milk (c), and mouse plasma (d), measured for 15 minutes at 0 (left), 15 (middle), and 40 (right) minutes after addition of EVs to each cell type.*

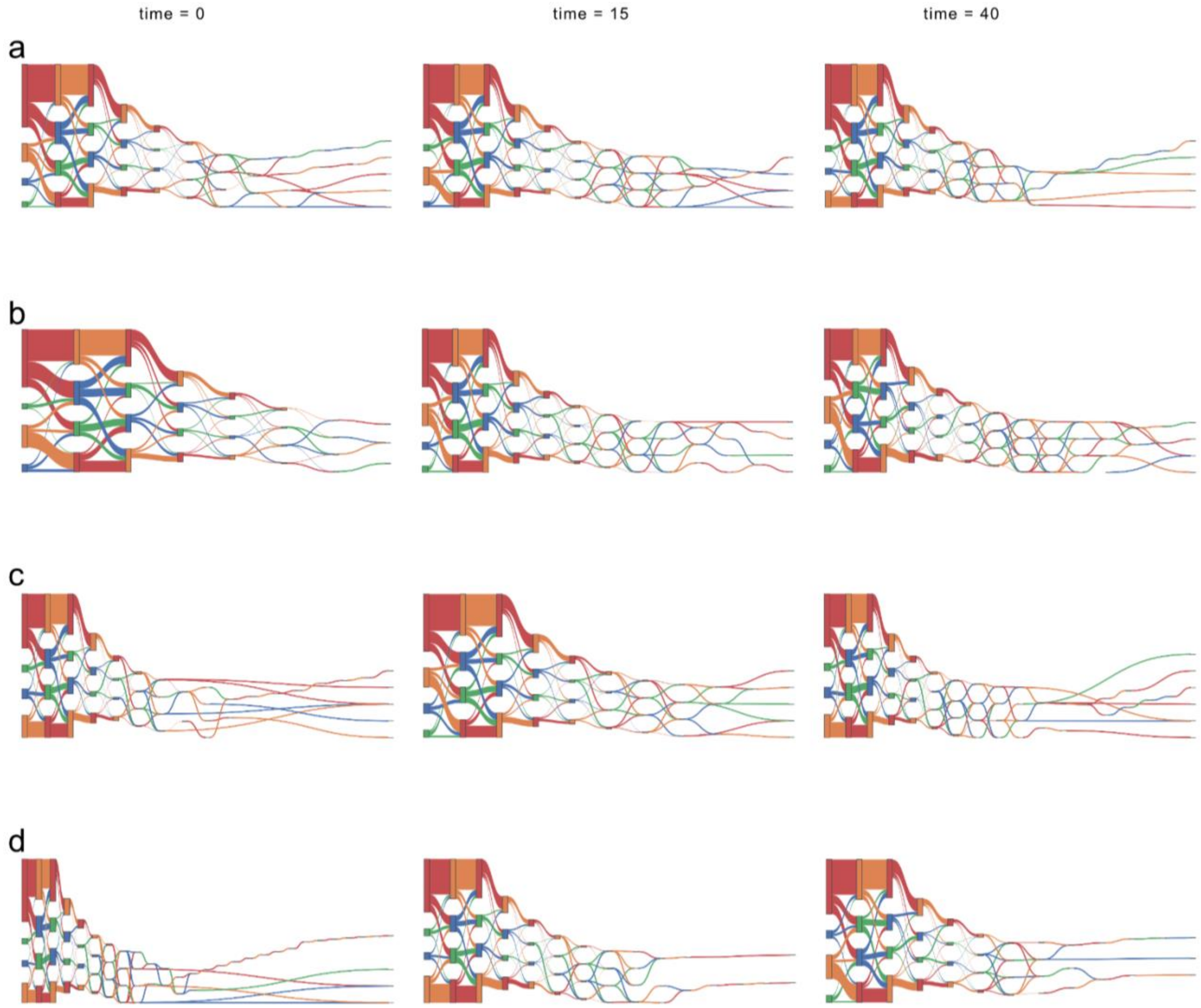

***SI Fig. 16: Sankey diagram of single-particle diffusive behavior over time.*** *Graph illustrating transitions between distinct diffusion modes along individual trajectories in HEK293-H cells. Extracellular vesicles from C2C12 cells (a), human plasma (b), whole cow milk (c), and mouse plasma (d), measured for 15 minutes at 0 (left), 15 (middle), and 40 (right) minutes after addition of EVs to each cell type.*

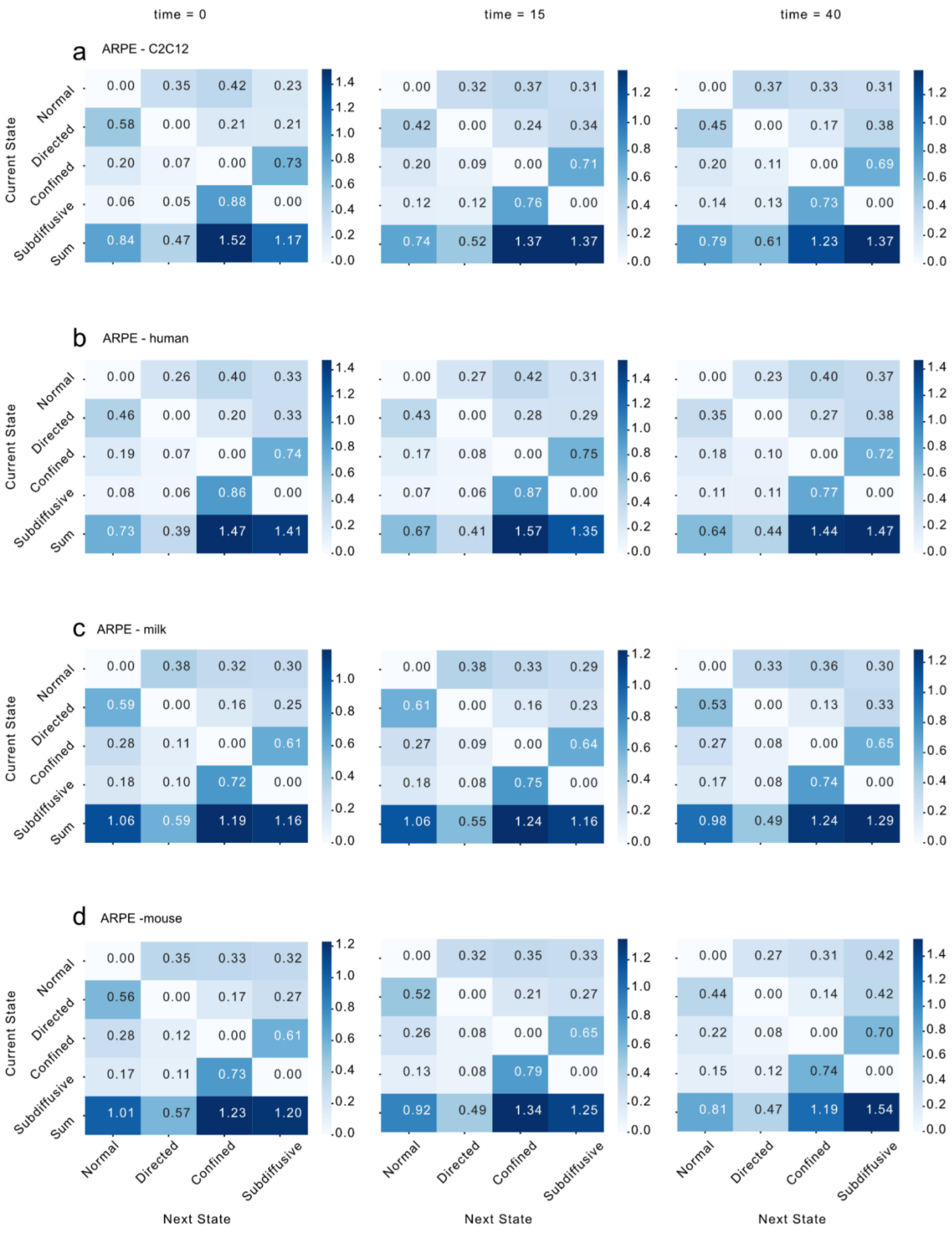
time = 0
time = 15
time = 40
a
ARPE - C2C12
b
ARPE - human
c
ARPE - milk
d
ARPE -mouse
Current State
Next State
Normal
Directed
Confined
Subdiffusive
Sum

***SI Fig. 17: Transition Probability Matrix from ARPE-19 cells.*** *Extracellular vesicles from C2C12 cells (a), human plasma (b), whole cow milk (c), and mouse plasma (d), measured for 15 minutes at 0 (left), 15 (middle), and 40 (right) minutes after addition of EVs to each cell type.*

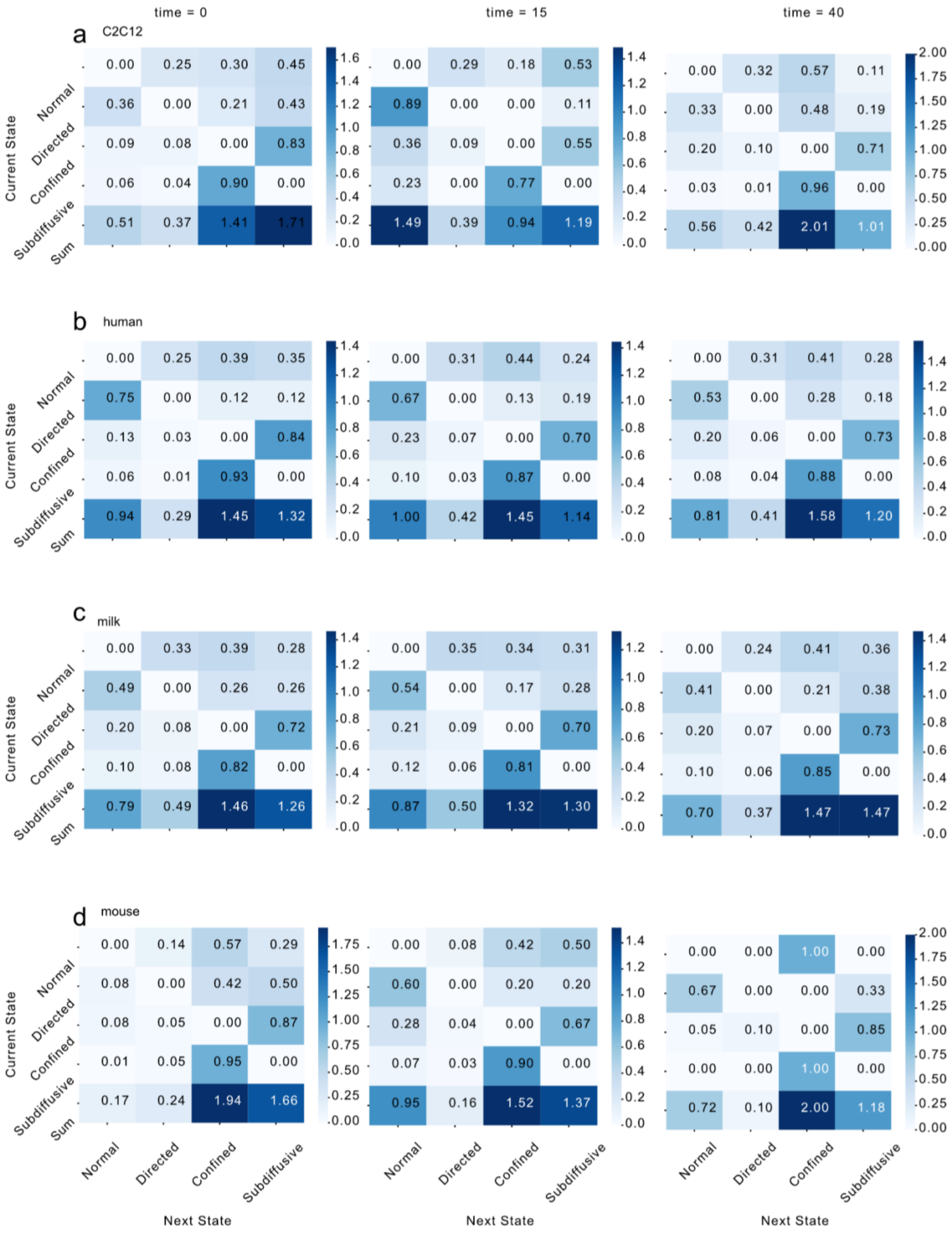
time = 0
time = 15
time = 40
a
C2C12
b
human
c
milk
d
mouse
Current State
Next State
Normal
Directed
Confined
Subdiffusive
Sum

***SI Fig. 18: Transition Probability Matrix from bEnd.3 cells.*** *Extracellular vesicles from C2C12 cells (a), human plasma (b), whole cow milk (c), and mouse plasma (d), measured for 15 minutes at 0 (left), 15 (middle), and 40 (right) minutes after addition of EVs to each cell type.*

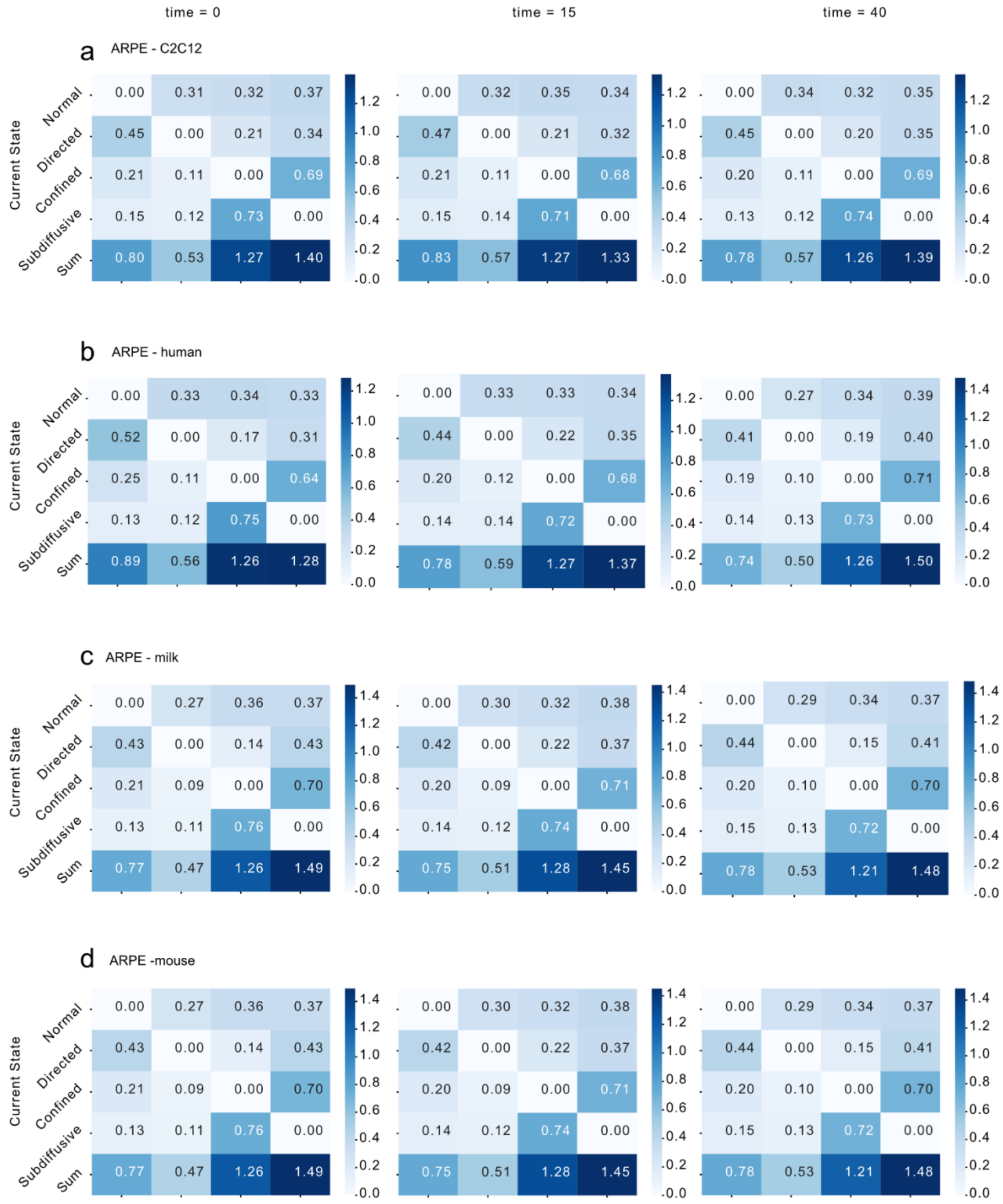

***SI Fig. 19: Transition Probability Matrix from HEK293-H cells.*** *Extracellular vesicles from C2C12 cells (a), human plasma (b), whole cow milk (c), and mouse plasma (d), measured for 15 minutes at 0 (left), 15 (middle), and 40 (right) minutes after addition of EVs to each cell type.*

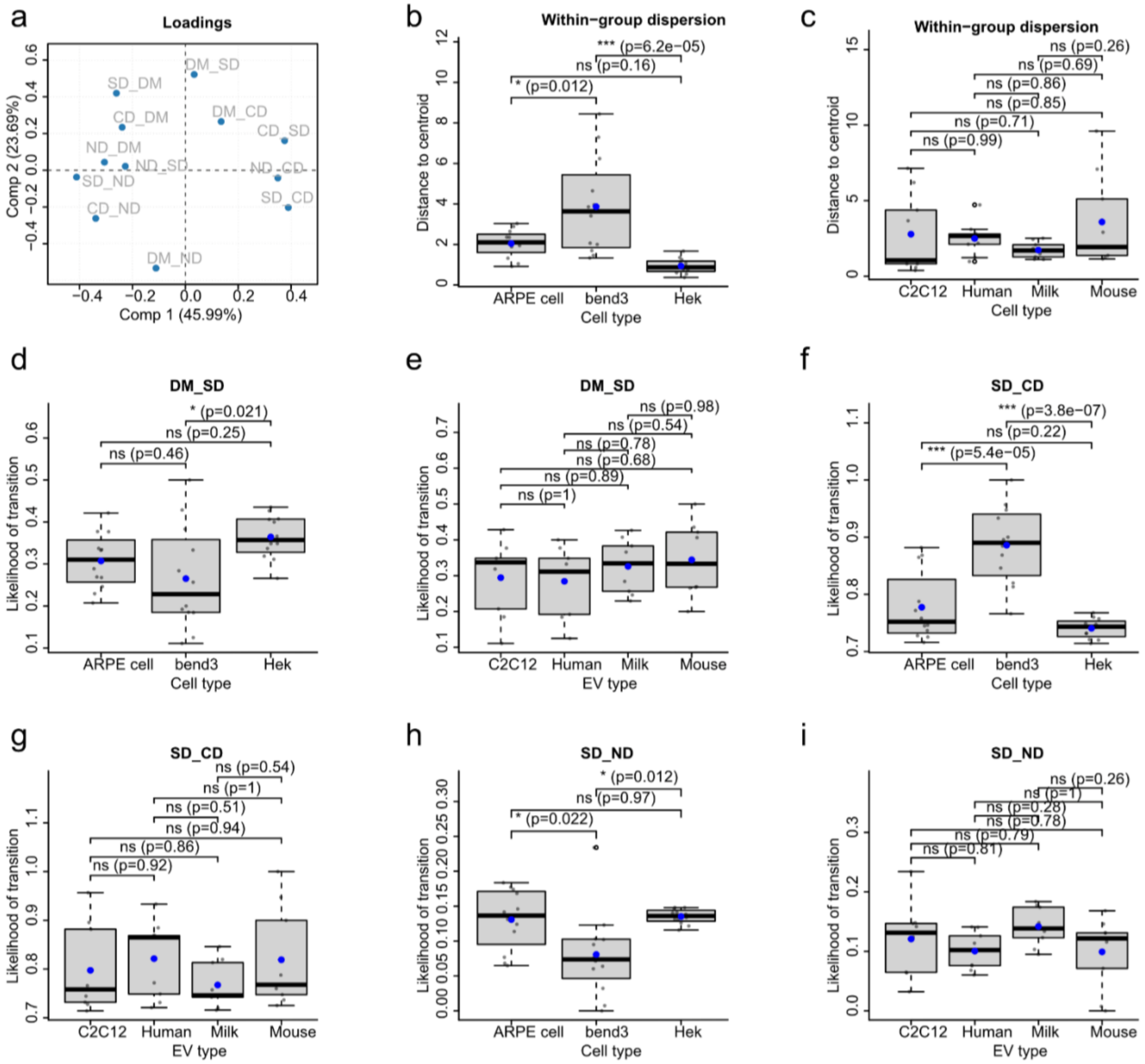

***SI Fig. 20: PCA analysis of transition probabilities.*** *(a) Principal component loadings (PC1 and PC2) highlighting variables contributing to variance in transition behavior.*
*(b–c) Within-group dispersion of EVs grouped by origin.*
*(d–i) Transition probabilities between diffusion states: (d,e) directed motion → subdiffusion, (f,g) subdiffusion → confined motion, and (h,i) subdiffusion → normal diffusion, grouped by recipient cell type (d,f,h) or EV origin (e,g,i). Asterisks denote statistical significance (*P ≤ 0.05, **P ≤ 0.01, ***P ≤ 0.001; ns, not significant, P ≥ 0.05), determined by one-way ANOVA with Tukey's post hoc test.*

**Data Availability**

All raw data are available upon request. Data supporting the findings of this study and Software code for treatment of single vesicle data and statistical analysis is available here: https://anon.erda.au.dk/cgi-sid/ls.py?share_id=ebd30fzJE3
